# Towards a dynamical understanding of microstate analysis of M/EEG data

**DOI:** 10.1101/2023.04.09.536154

**Authors:** Nikola Jajcay, Jaroslav Hlinka

**Author notes:** **For correspondence:** (NJ); (JH). National Institute of Mental Health, Topolová 748, Klecany, 250 67, Czechia. **Data availability:** Preprocessed public EEG LEMON dataset (Babayan et al., 2019); code: N.J.’s GitHub. **Funding:** NJ: HORIZON-WIDERA No. 101090306 (European Research Executive Agency); JH: Czech Science Foundation No. 21-32608S. **Competing interests:** The authors declare no competing interests.

## Abstract

One of the interesting aspects of EEG data is the presence of temporally stable and spatially coherent patterns of activity, known as microstates, which have been linked to various cognitive and clinical phenomena. However, there is still no general agreement on the interpretation of microstate analysis. Various clustering algorithms have been used for microstate computation, and multiple studies suggest that the microstate time series may provide insight into the neural activity of the brain in the resting state. This study addresses two gaps in the literature. Firstly, by applying several state-of-the-art microstate algorithms to a large dataset of EEG recordings, we aim to characterise and describe various microstate algorithms. We demonstrate and discuss why the three “classically” used algorithms ((T)AAHC and modified K-Means) yield virtually the same results, while HMM algorithm generates the most dissimilar results. Secondly, we aim to test the hypothesis that dynamical microstate properties might be, to a large extent, determined by the linear characteristics of the underlying EEG signal, in particular, by the cross-covariance and autocorrelation structure of the EEG data. To this end, we generated a Fourier transform surrogate of the EEG signal to compare microstate properties. Here, we found that these are largely similar, thus hinting that microstate properties depend to a very high degree on the linear covariance and autocorrelation structure of the underlying EEG data. Finally, we treated the EEG data as a vector autoregression process, estimated its parameters, and generated surrogate stationary and linear data from fitted VAR. We observed that such a linear model generates microstates highly comparable to those estimated from real EEG data, supporting the conclusion that a linear EEG model can help with the methodological and clinical interpretation of both static and dynamic human brain microstate properties.

## Introduction and Background

Electroencephalography (EEG) is a widely used neuroimaging technique for studying the electrical activity of the brain. One interesting aspect of EEG data is the presence of temporally stable and spatially coherent patterns of activity, known as microstates (Pascual-Marqui et al., 1995; Koenig et al., 1999; Koenig et al., 2002). Microstates have been found to reflect the underlying functional architecture of the brain and have been linked to various cognitive and clinical phenomena (Dietrich Lehmann et al., 2010; Schlegel et al., 2012; Kikuchi et al., 2007). Previous research on EEG microstates has used a variety of approaches for microstate derivation, including various clustering algorithms (Dietrich Lehmann et al., 1987; Pascual-Marqui et al., 1995; Khanna et al., 2014; Rukat et al., 2016). However, neither the relative performance nor the validity of these methods has not been systematically compared. The algorithms we compare in this study include classical microstate algorithms such as (Topographic) Atomize and Agglomerate Hierarchical Clustering ((T)AAHC) (Murray et al., 2008) and modified K-Means (Pascual-Marqui et al., 1995) and compare it with Principal Component Analysis decomposition (Haykin et al., 2004), Independent Component Analysis decomposition (Haykin et al., 2004) and Hidden Markov Models (Rezek et al., 2002).

In a seminal paper, Dietrich Lehmann et al., 1987 demonstrated that the alpha frequency band (8–12 Hz) of the multichannel resting-state EEG signal could be parsed into a limited number of distinct quasi-stable states. Since the introduction of microstate analysis, they have been termed as candidates for “atoms of thought” (Lehmann, 1993; Dietrich Lehmann et al., 1998) and used to describe the variability across behavioural states (Stevens et al., 1998; Dietrich Lehmann et al., 2010), personality types (Schlegel et al., 2012), and neuropsychiatric disorders (Dietrich Lehmann et al., 2005; Kikuchi et al., 2007). Some evidence suggests that the microstate time series may provide insight into the neural activity of the brain in the resting state by connecting fast and transient microstate switching to resting-state BOLD activity (Britz et al., 2010; Musso et al., 2010; Yuan et al., 2012). However, the functional interpretation and significance of microstate maps and how they relate to mental processes have not been established yet.

Moreover, to the best of our understanding, there is no general agreement on the validity of microstate analysis as such. In particular, we are still unsure whether the base assumption of microstate analysis, i.e., whether the brain dynamics actually switches between a relatively low and fixed number of discrete states, is indeed valid. To this end, numerous works studied the so-called low-dimensional manifolds of brain dynamics due to observations that the whole-brain imaging of spontaneous brain dynamics exhibits highly constrained patterns (Fox et al., 2007; Ponce-Alvarez et al., 2018), and therefore it has been hypothesized that the metastable dynamics govern brain activity (Tognoli et al., 2014; Deco et al., 2016; Rué-Queralt et al., 2021) and lies on a low-dimensional smooth manifold embedded in the high-dimensional space of the neuroimaging data (Jazayeri et al., 2017; Shine et al., 2019). However, this hypothesis assumes continuous attractors and smooth transitions (Gallego et al., 2017; Gallego et al., 2018; Chaudhuri et al., 2019), in stark contrast with the microstate assumption of discrete states and rapid switching.

Previous studies which explored the similarity of various clustering algorithms reported a high level of similarity between classical microstate clustering schemes and linear methods such as PCA. von Wegner et al., 2018 provided numerical and computational insights that multiple algorithms are information-theoretically invariant. Although some differences exist, these so-called static properties contain a significant amount of information about the clustering algorithm (e.g., decorrelation of PCA, which leads to slightly different topographies C and D (von Wegner et al., 2018)), the dynamic and information-theoretical properties reflect intrinsic dynamical properties of the EEG signal and are, therefore, independent of the employed algorithm (von Wegner et al., 2018). This finding leads us to hypothesise that dynamical microstate properties might be obtainable from linear characteristics of the underlying EEG signal. In particular, we might be able to estimate dynamical microstate properties from the cross-covariance and autocorrelation structure of the EEG data.

To this end, we aim to test our hypothesis in two steps. Firstly, by generating a Fourier transform surrogate EEG signal (Theiler et al., 1992) (a Monte Carlo technique prevalent in statistical testing) and comparing microstate properties in real and synthetic datasets. Secondly, we consider EEG data as a stationary Vector Autoregression (VAR) process. We will estimate its parameters, generate a stationary signal with the same linear characteristics as the original process (our EEG data), and study the stability and similarity of obtained microstate properties.

In summary, this study addresses two gaps in the literature. By applying several state-of-the-art microstate algorithms to a large dataset of EEG recordings, our goal is to evaluate the performance of these algorithms in terms of their ability to detect and characterize microstates accurately. By comparing various algorithms, we provide a valuable resource for researchers interested in using EEG microstates to gain insights into brain function. Secondly, we evaluate the hypothesis that dynamical microstate properties are directly obtainable from linear characteristics of underlying EEG data (cross-covariance and autocorrelation). This would implicate the possibility of devising more efficient estimates, non-discrete generalizations, better explainability, and even the analytical treatment of microstate properties, which would be beneficial for relating various clinical findings from microstate literature to the brain dynamics (dys)function. Overall, this study represents a significant theoretical contribution to the field of EEG microstate research.

## Methods & Materials

### Microstate Analysis

#### Overview

Within microstate analysis, the multichannel EEG signal is viewed as a series of instantaneous topographies of electrical potential, encoded as a 2D matrix *X*_*ij*_ of shape (*n*_*ch*_, *n*_*t*_), with *n*_*ch*_ representing the number of channels and *n*_*t*_ the number of time points. Before clustering the signal, we extracted the global field power (GFP hereafter) peaks to account only for temporal points with the highest signal-to-noise ratio, as the EEG topographies are most clearly defined at these maxima, according to the standard procedure (Dietrich Lehmann et al., 1980; Dietrich Lehmann et al., 1987; Koenig et al., 1999). The GFP at each temporal instant is equal to the root mean square across the average-referenced potentials (i.e., a standard deviation of the signal across space):

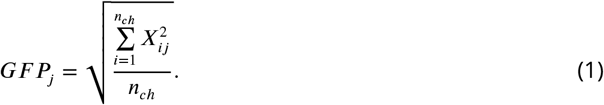

Note that in the microstate analysis, all EEG data are supposed to be average-referenced; therefore, we automatically have **∑**_*i*_ ***X***_*ij*_ = 0 for all *j*. Potential topographies (or maps) that occur at maxima of GFP time series represent instants of the highest field strength and greatest signal-to-noise ratio. The microstate analysis then includes two steps: the first is to identify a set of representative microstate maps, and the second is to project the original multichannel EEG recording into the basis comprised of these microstate maps, thus converting the EEG signal into a sequence of discrete microstate maps (see Fig. 1 for an overview).

**Figure 1.**
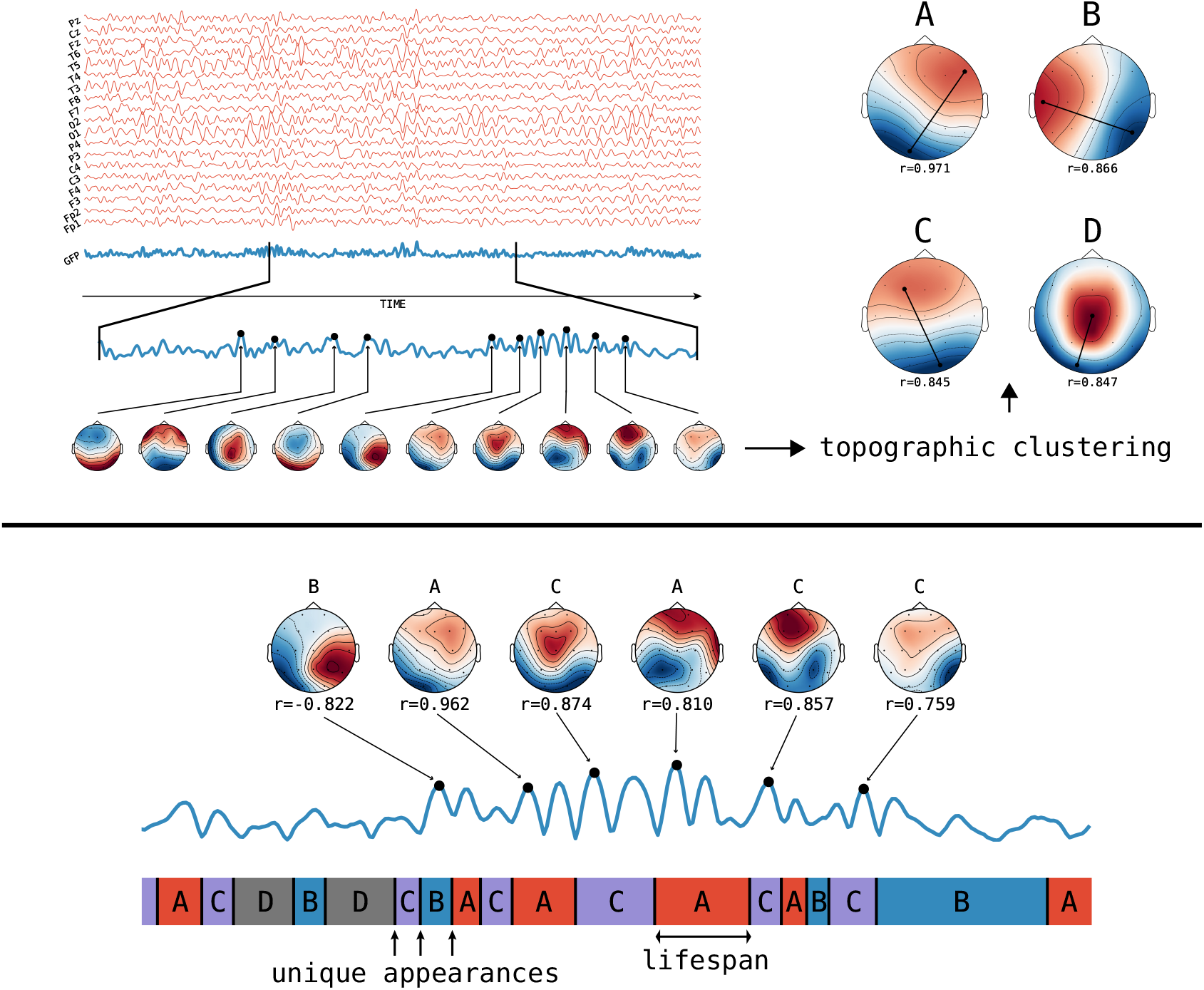
Schematic of the method of microstate analysis and extraction of features of interest from the microstate time series. The figure is adapted from Khanna et al., 2014. The GFP (in blue) is calculated at each instant of the multichannel EEG recording (various channels in red). Peaks of the GFP curve represent moments of the highest signal-to-noise ratio. At peaks of the GFP curve, the potential recorded at each electrode of the multichannel signal is plotted onto a map of the channel array. This collection of maps is entered into a microstate algorithm (topographic clustering), yielding a small number of canonical maps (**the microstates**) that explain a large proportion of the global topographic variance. Four topographies are repeatedly found using this method; these maps are labelled A, B, C, or D in the figure. Connected dots indicate points of maximum or minimum recorded electric potential. Then, the original maps at peaks of the GFP curve are assigned to a microstate class A, B, C, or D based on the degree of correlation with the microstate maps. This reassignment represents the original multichannel data as an alternating series of microstates A, B, C, and D. A microstate is considered dominant in the time during which all successive original maps are assigned to the same microstate class. Each period of dominance is considered a unique appearance of a microstate. The frequency of a microstate is the number of unique appearances per second. The coverage of each microstate is the fraction of the total recording time that each microstate is dominant.

Each algorithm thus yields a set of ***M*** microstate maps representing the EEG data set. After finding representative microstate maps, the original multichannel EEG recording was transformed into a temporal sequence of microstate maps by the following procedure: one of the canonical maps is assigned for each GFP peak as per the highest Pearson product-moment correlation. After this assignment of microstate identities to the GFP peaks, microstate identity is further rolled out to all other time points according to the microstate identity of the temporally closest GFP peak.

The computation of microstate sequences is identical for all clustering methods, following a “winner takes all” approach, also called competitive back-fitting (Pascual-Marqui et al., 1995). To measure how well the microstate sequence approximates the underlying EEG data set, a frequently used quantity is global explained variance (GEV) (Murray et al., 2008). GEV measures the percentage of data variance explained by given microstate maps.

All clustering algorithms used in our study can yield a different number of microstate maps, as the number of clusters is actually a parameter of the method(s). For better compatibility and to avoid unnecessary complexity, we chose to use a fixed number of four microstates for all algorithms, in line with seminal early works in this area (Dietrich Lehmann et al., 1987; Pascual-Marqui et al., 1995) and with new studies (Rieger et al., 2016; von Wegner et al., 2018). The resulting microstate maps are ordered according to the template maps provided by Koenig et al. (Koenig et al., 2002) to ensure replicability and comparison between algorithms.

#### Clustering Algorithms

In our work, we compared 6 different algorithms that can be used for the clustering stage of the microstate analysis:

##### (Topographic) Atomize and Agglomerate Hierarchical Clustering

The Atomize and Agglomerate Hierarchical Clustering (AAHC) and the related Topographic AAHC (TAAHC) are deterministic hierarchical clustering algorithms. Both algorithms use a bottom-up approach where they are initialised with each EEG topography being a separate cluster, and they iteratively reduce the number of clusters. At each iteration, the worst cluster is atomised, and all its former members are reassigned to a different cluster one by one (Murray et al., 2008; Khanna et al., 2014). The algorithms differ in how the worst cluster is determined: in the AAHC, the worst cluster is simply the one with the lowest GEV, while in the TAAHC algorithm, the worst cluster is the one with the lowest intra-cluster similarity. The reassignment of former worst cluster members is done via maximum similarity. The algorithm concludes when the pre-selected number of clusters is achieved.

##### Modified K-Means

The modified K-means algorithm is historically the first algorithm used for microstate analysis (Pascual-Marqui et al., 1995). It is an iterative stochastic clustering algorithm inspired by classical K-means clustering (Lloyd, 1982). The algorithm is initialised with randomly selected GFP maxima topographies as cluster centres. Then it proceeds in a two-step iterative fashion by estimating the microstate sequence using current cluster centres and updating the cluster centres based on the sequence. The iteration continues until convergence, where the cluster centres no longer change. Due to the stochasticity of algorithm initialisation of the algorithm, different runs might yield different resulting microstate maps and microstate assignments. Typically, this whole procedure is thus run multiple times (e.g., 100 times), and the best run according to the GEV is selected.

##### Principal Component Analysis

Principal Component Analysis (PCA) is one of the most frequently used methods for dimensionality reduction due to its simplicity and straightforward interpretation (Haykin et al., 2004). Spatial PCA has previously been used to cluster EEG topographies (Skrandies, 1989; Spencer et al., 1999; Spencer et al., 2001). Here, PCA is computed by eigendecomposition of data covariance matrix 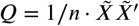, where 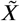 contains the multichannel EEG signal at GFP peaks and the prime denotes matrix transposition. The principal components (i.e., the canonical microstates or the cluster centres) correspond to the eigenvectors of the matrix *Q*. The PCA naturally orders the components regarding their respective eigenvalues, and the components are mutually orthogonal. Like the (T)AAHC algorithm, PCA is deterministic and naturally fosters reproducibility.

##### Independent Component Analysis

Independent Component Analysis (ICA) is another widely used decomposition technique in neuroscientific applications (Haykin et al., 2004). ICA seeks statistical independence between components, i.e. ICA performs so-called signal unmixing. Signal unmixing is typically written as 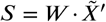, where 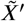 contains the transposed matrix of the EEG data at GFP peaks, *W* denotes the unmixing matrix, and *S* is the so-called source matrix. The particular implementation we relied on is Fast-ICA (Hyvärinen et al., 2000), with a log-cosh activation function and pre-whitening.

##### Hidden Markov Model

In contrast to other clustering algorithms used in our study, the Hidden Markov Model (HMM) is a generative model that describes observations emerging from the rapid switching of hidden states with a Gaussian emission model (Rezek et al., 2002). HMM was successfully used to analyse resting-state MEG data and compare them to classical resting-state networks as studied in fMRI (Baker et al., 2014). Here, we assume an HMM of length *n*_*t*_ samples, state space dimension of 4 (as per the number of latent states), hidden state variables 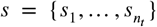 and observed data *X*_*it*_ with *t* ∈ [0.. *n*_*t*_]. The full posterior probability of our HMM is then (Baker et al., 2014):

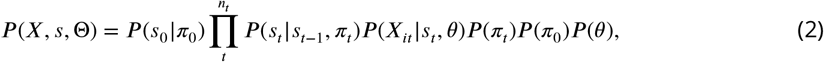

where *P*(*π*)_*t*_), *P*(*π*_0_), and *P*(*θ*) are chosen as non-informative priors. The parameters of HMM Θ = {*π*_0_, *π*_*t*_, *θ*}, subject to fitting, consist of *π*_0_, which parametrises the initial state probability *P*(*s*_0_), *π*_*t*_ which dictates the state transitions *P*(*s*_*t*_ |*s*_*t*−1_), and *θ* which describes the observation probability *P*(*X*_*it*_|*s*_*t*_). We assume that the model is Markovian, i.e., that the probability of transition to another state depends only on the state that the system is in. The term *P*(*X*_*it*_|*s*_*t*_, *θ*) describes the observation (emission) model, and we assume that this emission model is a multivariate normal distribution with

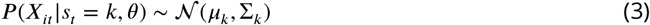

for state *k*. Our implementation uses variational Bayes inference on our HMM, with Viterbi decoding yielding the inference on state variables *P*(*s*_*t*_|*X*_*it*_), i.e., the sequence of hidden states.

#### Microstate Measures

In our study, we use a number of summary statistics to describe the temporal characteristics of inferred sequences from various clustering algorithms. For the computation of statistics, we use a sequence of latent states, *u*_*t*_, obtained directly from our algorithms (e.g. for ICA and PCA, the maximum activation in each time step, for HMM the Viterbi decoding, etc.), albeit the sequences were relabelled such that the microstate topographies match the Koenig et al., 2002 templates. This also ensures comparability among subjects and algorithms, e.g., state A corresponds to the same literature-based template microstate (Koenig et al., 2002) for all subjects and all clustering algorithms.

##### Average lifespan

The lifespan of microstates is computed as the time during which all successive original data points were assigned the same microstate class *k* (Dietrich Lehmann et al., 2005) (cf. Fig. 1B), i.e.

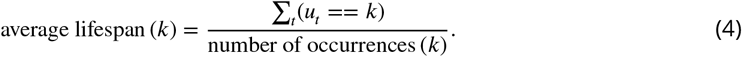

##### Coverage

The coverage of the microstate is calculated by taking a ratio of the total time spent in a particular microstate *k* over the total recording time (Dietrich Lehmann et al., 2005):

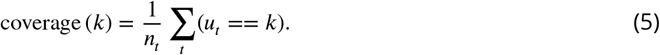

##### Occurrence

Microstate frequency is determined by counting unique appearances of a microstate *k* in one second of the recording (Dietrich Lehmann et al., 2005):

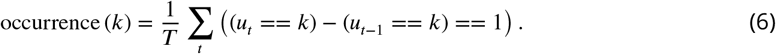

### Synthetic Data

For the theoretical part of our study, we generated synthetic data by sampling from a multivariate normal distribution, *X*_*i,j*_ ∼ **𝒩**(*μ*, **∑**), with a zero mean vector (*μ*_*i*_ = 0 for *i* ∈ [0.. *n*_*ch*_]) and a random covariance structure with **∑** = ***M*·*M***′ where ***M***_*ij*_ ∼ **𝒩**(0, 1) for *i, j* ∈ [0.. *n*_*ch*_]. To mimic the microstate analysis pipeline, we used “average reference” the data, i.e., normalise to zero mean over channels for each time point separately. For various experiments, we varied the number of channels *n*_*ch*_ and sampled 2500 time points, mimicking a 10-second long time series with a sampling frequency of 250 Hz.

### Experimental Data

For the experimental part of our study, we used publicly available EEG data that are part of the Max Planck Institut Leipzig Mind-Brain-Body Dataset (LEMON) dataset (Babayan et al., 2019). The dataset consists of 228 healthy participants comprising a young (*N* = 154, 25.1 ± 3.1 years, range 20–35 years, 45 female) and an elderly group (N=74, 67.6±4.7 years, range 59–77 years, 37 female) acquired cross-sectionally in Leipzig, Germany, between 2013 and 2015. Among other neuroimaging modalities, participants completed a 62-channel EEG experiment at rest (rsEEG hereafter) using two paradigms: eyes open and eyes closed. We used directly the preprocessed EEG data (total *N* = 204) provided as EEGLAB `.set` and `.fdt` files. The complete data description and EEG pre-processing pipelines can be found elsewhere (Babayan et al., 2019). Briefly, all EEG data have a sampling frequency of 250 Hz, are low-pass-filtered with a 125 Hz cutoff frequency, and are approximately 8 minutes long.

### Surrogate Data

In order to evaluate the microstate analysis pipeline on random data resembling the real EEG data, we used the bootstrapping technique known as surrogate data (Paluš, 2007). The idea is to preserve some statistics of the data while randomizing others. In our study, we used multivariate Fourier Transform surrogates (Theiler et al., 1992) (also called random phases), which preserve spectral content and autocorrelation function, but destroy any tentative relationships between temporal scales. The surrogate procedure used in our study yields time series data of the same length and sampling rate as the original EEG data. We used a fixed seed (42) for a random number generator, such that we are able to compare surrogate procedures across participants.

### Vector Autoregression Data

To test our hypothesis that microstate properties represent and can be estimated from linear covariance and autocovariance structure, we estimate a Vector Autoregression process VAR(*p*) of order *p*, approximating the experimental EEG data. The VAR(*p*) models a *T* × *K* multivariate time series *Y*, where *T* denotes the number of time steps (observations), and *K* denotes the number of variables (EEG channels), as

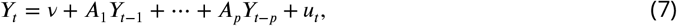

with *u*_*t*_ ∼ 𝒩(0, ∑_*u*_), and *A*_*i*_ the *K* × *K* coefficient matrices (Lütkepohl, 2005).

We start by estimating the order *p* of the VAR process for each subject in our dataset. Here we fit VAR(*p*) for each subject separately with *p* ∈ [0, …, 10] and select the order which minimises the Akaike Information Criterion (AIC). We then simulate VAR(*p*) for each subject for 1200 or 3600 seconds (with the same sampling rate as original EEG data) with random initialisation. Finally, we run our microstate analysis in two branches: the first branch estimates microstates and their properties on full, long VAR data, and the second branch estimates microstates and their properties on segments of varying lengths (10, 30, 60, or 180 seconds) and then averages properties over these segments.

### Computational methods and implementation

We implemented all our analyses, including clustering algorithms, all measures and statistical tests in `python` version 3.7. For the loading and processing of experimental data, we relied on the `mne` library (Gramfort et al., 2013). For ICA and PCA clustering algorithms, we used the implementation in the `scikit-learn` library (Pedregosa et al., 2011). For Hidden Markov Model, we used the `hmmlearn` library, and for other clustering algorithms, we wrote our custom implementation. The whole codebase can be found on N.J.’s GitHub. Together with the publicly available experimental dataset (cf. section Experimental Data) that we used in this study, we ultimately adhere to the FAIR principles for scientific data management (Wilkinson et al., 2016), and therefore all our results are directly replicable. Moreover, we registered our project on the Open Science Foundation (OSF) registry and uploaded our results on `figshare` platform connected to our OSF project.

## Results

### Understanding Clustering Algorithms on Synthetic Data

This section evaluates all 6 clustering algorithms on simple synthetically generated data. For ease of visualization, we opted to generate 3D multivariate normal data with average referencing, mimicking a 3-channel EEG montage with a sampling rate of 250 Hz. We then run the clustering algorithm in two ways: using only the GFP peaks (as is usual in the microstate analysis pipeline) or on a whole time series to evaluate the effect of this transformation. As will be apparent in later figures, selecting GFP peaks corresponds to the non-equidistant subsampling in the temporal sense and selecting values further away from the mean in the phase space, i.e., the edges of the multivariate data cloud in *n*_*ch*_ dimensions.

#### (Topographic) Atomize and Agglomerate Hierarchical Clustering

The most used classical microstate algorithms are the Atomize and Agglomerate Hierarchical Clustering and its Topographic version. Both are hierarchical, which means they proceed bottom-up in the sense that they are both initialised with each EEG topography as a separate cluster, and these are then iteratively merged to form the final 4 clusters. The result of applying (T)AAHC algorithms is illustrated in Figs. 2 and 3, respectively.

**Figure 2.**
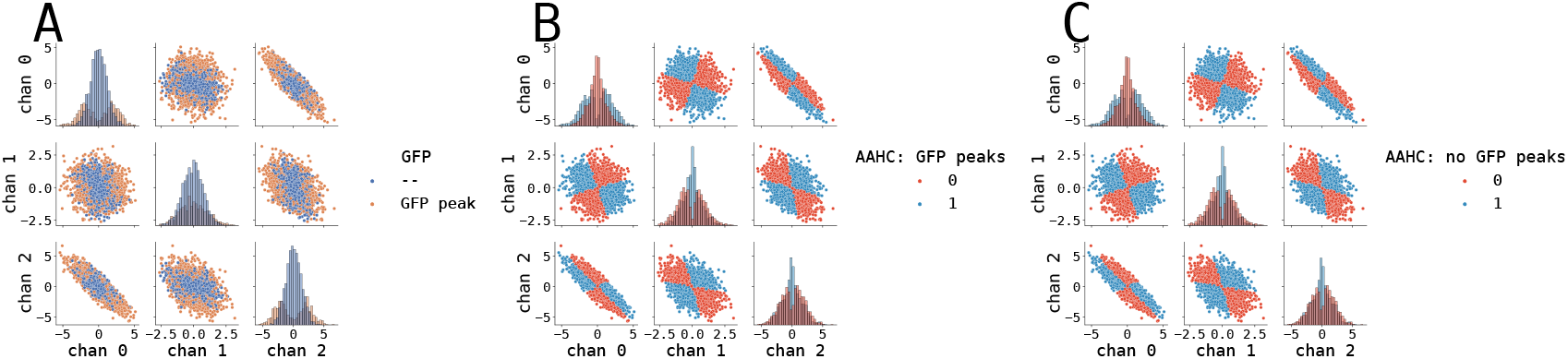
AAHC clustering algorithm result on synthetic data. We generated a synthetic 3D dataset as a multivariate normal distribution of length 10 seconds with a 250Hz sampling rate (cf. section Synthetic Data). (**A**)The data distribution, where GFP peaks are coloured in orange. Note how GFP peaks are located on the edge of the data cloud. (**B**) Application of AAHC algorithm on synthetic data using only GFP peaks. (**C**) Application of AAHC algorithm on synthetic data using all available data points.

**Figure 3.**
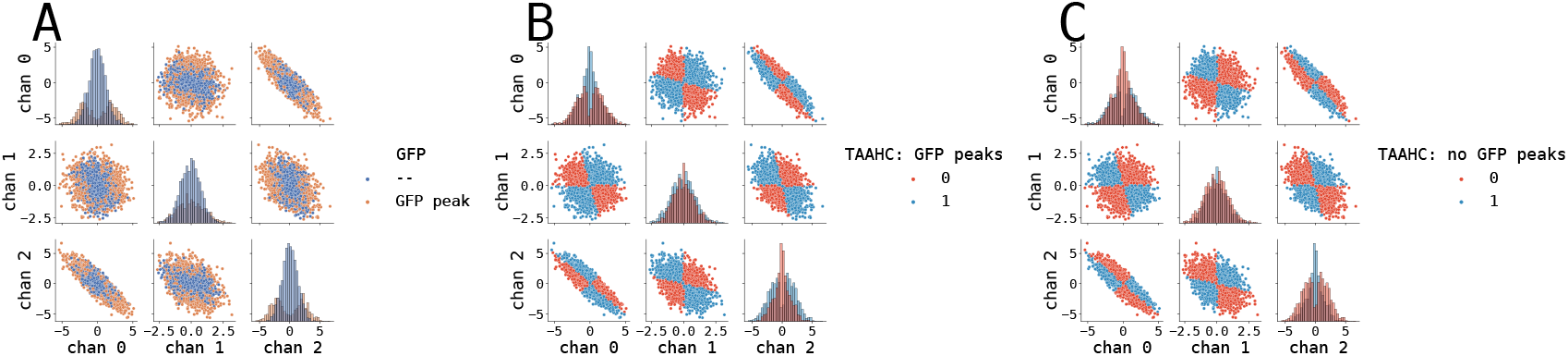
TAAHC clustering algorithm result on synthetic data. We generated a synthetic 3D dataset as a multivariate normal distribution of length 10 seconds with a 250Hz sampling rate (cf. section Synthetic Data). (**A**)The data distribution, where GFP peaks are coloured in orange. Note how GFP peaks are located on the edge of the data cloud. (**B**) Application of TAAHC algorithm on synthetic data using only GFP peaks. (**C**) Application of TAAHC algorithm on synthetic data using all available data points.

Both bottom-up algorithms show very similar segmentation results for both cases of preprocessing: using only GFP peaks or the whole synthetic time series. The reason why GFP preprocessing does not change the segmentation results is simple: since both algorithms work in the bottom-up sense, they purely agglomerate the most similar clusters together (AAHC and TAAHC only differ in the agglomeration criterion). In the case of the AAHC algorithm, the topographies that are not GFP peaks (and, thus, have low variance (GEV)) are merged into clusters with higher GEV (hence, the topographies that are GFP peaks). Then, as the cluster centroids are in (T)AAHC defined by the first eigenvector, these low-GEV maps have only negligible effect on the resulting centroids. A similar argument is valid for TAAHC. To quantify the similarity effect, we also computed similarity measures between the segmentations of only GFP peaks run and whole data run, using Spearman correlation (**r**) entailed with analytical p-value and accuracy (**acc**) that was statistically assessed using a permutation-based p-value. Naturally, when the algorithm switched polarity (swapped classes, i.e., 0 → 1 and 1 → 0), we manually swapped cluster assignments. For the AAHC algorithm, the similarity reached **r**(2500) = 0.995, *p* < 0.001 and **acc** = 0.998, *p* < 0.001 (20000 permutations), while for TAAHC algorithm the similarity reached **r**(2500) = 0.993, *p* < 0.001 and **acc** = 0.996, *p* < 0.001 (20000 permutations).

#### Modified K-Means

The modified K-Means algorithm is the original classically used clustering algorithm in the microstate analyses. The result of applying the modified K-Means algorithm on our synthetic dataset to find 2 microstate maps is shown in Fig. 4.

**Figure 4.**
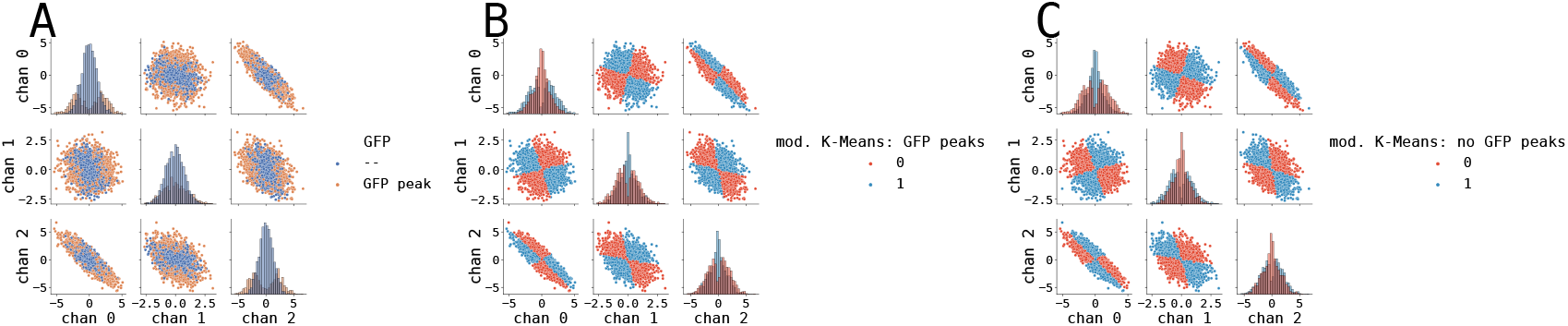
Modified K-Means clustering algorithm result on synthetic data. We generated a synthetic 3D dataset as a multivariate normal distribution of length 10 seconds with a 250Hz sampling rate (cf. section Synthetic Data). (**A**)The data distribution, where GFP peaks are coloured in orange. Note how GFP peaks are located on the edge of the data cloud. (**B**) Application of modified K-Means algorithm on synthetic data using only GFP peaks. (**C**) Application of modified K-Means algorithm on synthetic data using all available data points.

In the case of modified K-Means clustering, the preprocessing when we use only the peaks of the GFP curve does not change the segmentation much. This is obvious when we look inside the algorithm and interpret it geometrically. The algorithm proceeds iteratively in two steps: firstly, it computes the assignments of input topographies by projecting the input vector onto cluster centres and finding the direction with maximum loading. Then, it recomputes cluster centroids as a weighted average over topographies of that cluster, weighted by the Euclidean norm of a particular map. This distance weighting will naturally skew the final centroids towards topographies further away, which, coincidentally, are also the maps selected by the GFP peak procedure. Both similarity measures thus show very high similarity between the GFP and no GFP segmentations (**r**(2500) = .945, *p* < 0.001, and **acc** = 0.972, *p* < 0.001 (20000 permutations)).

#### Hidden Markov Model

The hidden Markov model is a generative model that describes the observations that emerge from rapid switching between topographies with a Gaussian observation model. To the best of our knowledge, it is not typically used on par with microstate analysis, albeit it promises to capture temporal and spatial dynamics that are more closely related to underlying brain activity (using the hidden states) than classical microstate analysis (Rukat et al., 2016). The result of applying the hidden Markov Model on our synthetic dataset is shown in Fig. 5.

**Figure 5.**
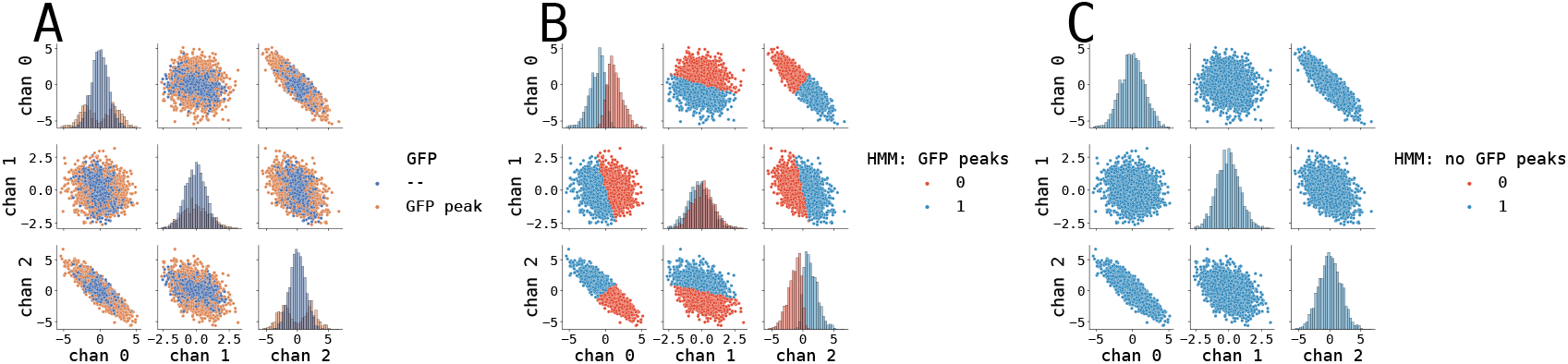
Hidden Markov Model clustering algorithm result on synthetic data. We generated a synthetic 3D dataset as a multivariate normal distribution of length 10 seconds with a 250Hz sampling rate (cf. section Synthetic Data). (**A**)The data distribution, where GFP peaks are coloured in orange. Note how GFP peaks are located on the edge of the data cloud. (**B**) Application of hidden Markov model algorithm on synthetic data using only GFP peaks. (**C**) Application of hidden Markov model algorithm on synthetic data using all available data points.

The segmentation results for GFP and no GFP case for HMM algorithm differ significantly. This is mainly given by the observation model in the HMM, which is Gaussian. When we use GFP preprocessing, we select points further away from the centre (see Fig. 5A). Given that the data does not exhibit the same covariance in all dimensions, this also means the data are effectively “cut” the cloud into halves (at least in one dimension) and therefore the HMM algorithm will divide the cloud into two clusters in such a way, that both clusters are as close as possible to multivariate Gaussians (see histograms on the diagonal in Fig. 5B). When we do not use GFP peaks preprocessing, the HMM algorithm estimates correctly that the whole data cloud is one multivariate Gaussian. It thus assigns the same cluster to all points (Fig. 5C). Naturally, the similarity between these two approaches is very low (**r**(2500) = 0.021, *p* = 0.298, **acc** = 0.520, *p* = 0.480 (20000 permutations)).

#### Principal Component Analysis

Although not strictly a microstate algorithm, PCA is one of the most frequently used dimensionality reduction algorithms. Spatial PCA has been used to cluster EEG topographies mainly in the context of ERP experiments (Spencer et al., 1999; Spencer et al., 2001). In contrast to classical microstate algorithms such as modified K-Means, or (T)AAHC, PCA is deterministic and hence valuable when reproducibility is essential. Fig. 6 shows the segmentation given by PCA on our synthetic dataset.

**Figure 6.**
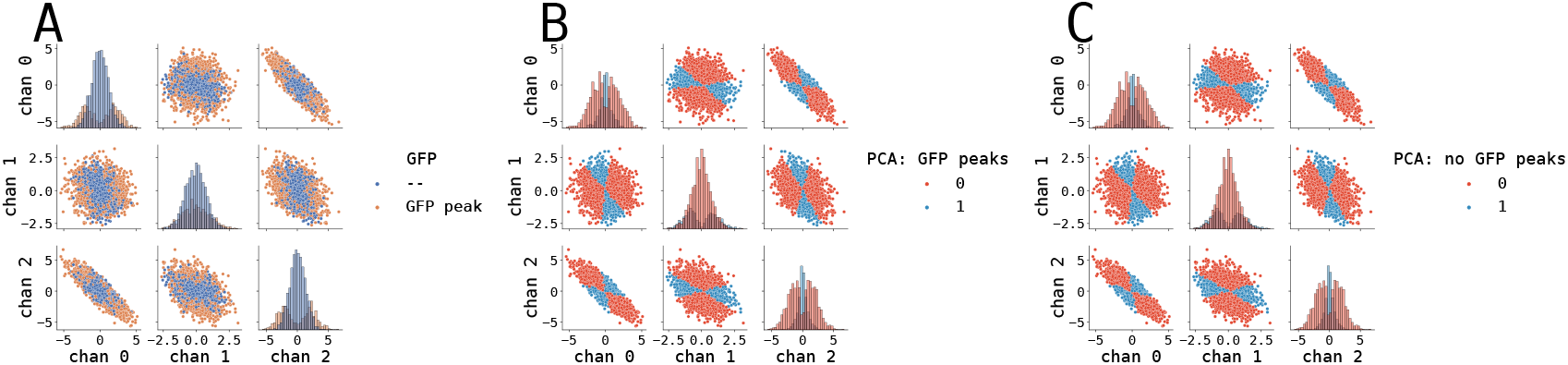
Principal component analysis clustering algorithm result on synthetic data. We generated a synthetic 3D dataset as a multivariate normal distribution of length 10 seconds with a 250Hz sampling rate (cf. section Synthetic Data). (**A**)The data distribution, where GFP peaks are coloured in orange. Note how GFP peaks are located on the edge of the data cloud. (**B**) Application of Principal component analysis algorithm on synthetic data using only GFP peaks. (**C**) Application of Principal component analysis algorithm on synthetic data using all available data points.

The results for PCA clustering look similar whether we use GFP peaks preprocessing or not. The reason is simple: PCA always selects the first principal component as the one with the highest variance, i.e., it follows the elongation as seen in the dataset (e.g., Fig. 6B top right). Removing points from the low variance region (i.e., the blue data points in the middle in Fig. 6A) does not change the shape of the remaining data. As the shape stays the same, so does the variance in all directions, and therefore the PCA always selects the same principal component. Indeed, the quantified similarity is very high: **r**(2500) = 0.987, *p* < 0.001 and **acc** = 0.995, *p* < 0.001 (20000 permutations).

#### Independent Component Analysis

Similarly to PCA, ICA is also a widely used technique in analysing neuroimaging data sets, particularly fMRI, where it is typically used to estimate so-called resting-state networks from resting-state fMRI data (Kiviniemi et al., 2003; Beckmann et al., 2005). Where PCA is based on spatial decorrelation, ICA seeks statistically independent components. However, the main criterion of independence may not necessarily correspond to biological relevance (for instance, statistically dependent EEG topographies might not be revealed by ICA) (for instance, statistically dependent EEG topographies might not be revealed by ICA) (von Wegner et al., 2018). The result of ICA clustering on our synthetic dataset is shown in Fig. 7.

**Figure 7.**
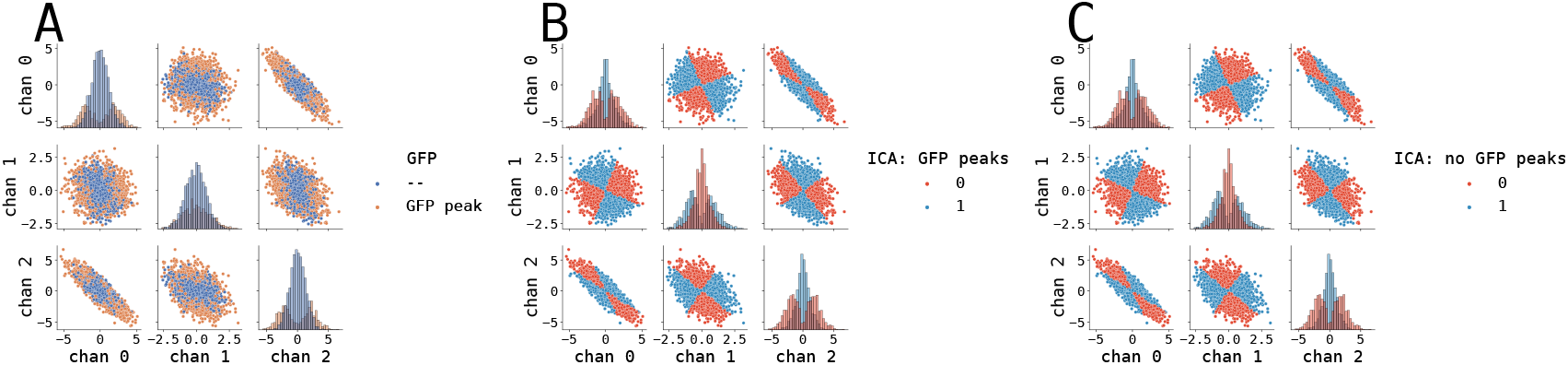
Independent component analysis clustering algorithm result on synthetic data. We generated a synthetic 3D dataset as a multivariate normal distribution of length 10 seconds with a 250Hz sampling rate (cf. section Synthetic Data). (**A**)The data distribution, where GFP peaks are coloured in orange. Note how GFP peaks are located on the edge of the data cloud. (**B**) Application of Independent component analysis algorithm on synthetic data using only GFP peaks. (**C**) Application of Independent component analysis algorithm on synthetic data using all available data points.

As with the PCA, ICA clustering yields very similar segmentation for both cases of preprocessing. Similarity measures are also very high: **r**(2500) = 0.997, *p* < 0.001 and **acc** = 0.998, *p* < 0.001 (20000 permutations).

Overall, we can conclude that, except for the Hidden Markov Model, all tested algorithms yield similar segmentations on our synthetic dataset in both tested cases: preprocessing with GFP peaks selection and without. The only argument for selecting GFP peaks would be computation speed, particularly for bottom-up agglomerative (T)AAHC algorithms. The hidden Markov Model is a special case here: selecting GFP peaks somewhat divides our dataset into two multivariate Gaussians, which is immediately picked up by the HMM, whereas keeping the full synthetic dataset (which was created as a multivariate Gaussian) leads to all points being assigned to the same cluster.

#### Synthetic Data Microstate Properties

In order to compare the six studied algorithms, we computed the overall similarity of their topographies (i.e., cluster centres) and segmentations for the 3D data presented above and compared their properties. The grand comparison is shown in Fig. 8.

**Figure 8.**
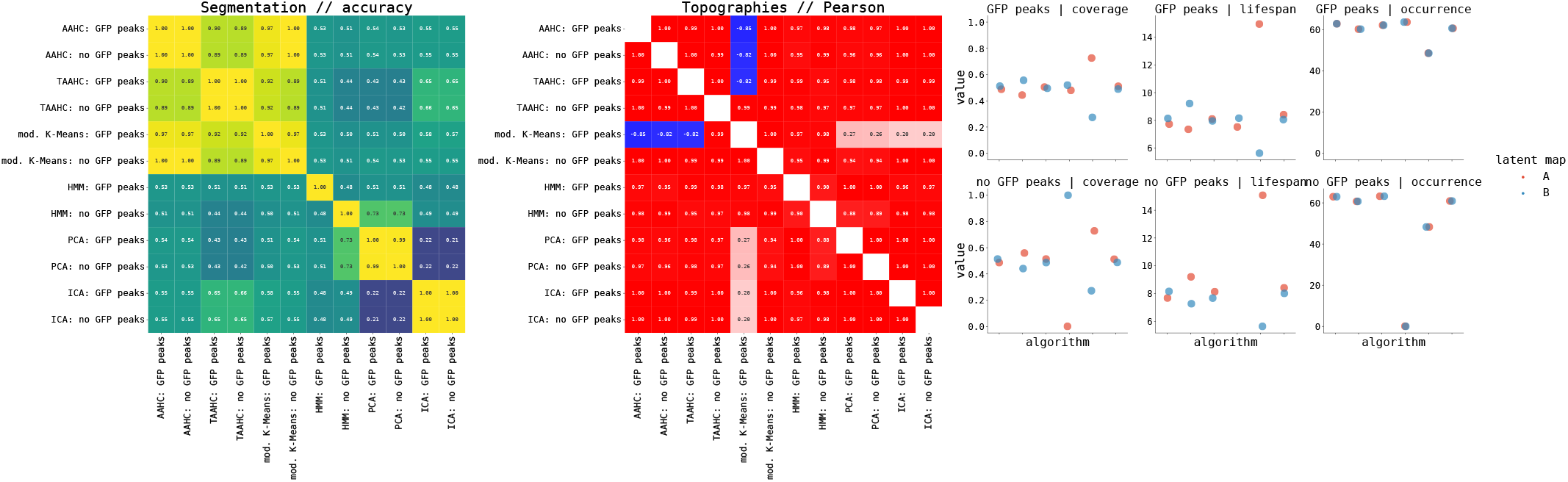
Microstate properties on the synthetic dataset using 6 algorithms for latent decomposition. (left) The similarity of segmentation as computed using 6 different microstate algorithms in 2 preprocessing options each (GFP or no GFP) is shown in the top left corner. The measure used to quantify similarity was accuracy. (middle) The similarity of topographies (cluster centres) yielded by each algorithm is shown in the bottom left corner using Pearson correlation. (right) Microstate properties; top row shows coverage, lifespan, and occurrence for GFP peaks preprocessing (cf. section Microstate Measures), and the bottom row shows coverage, lifespan, and occurrence for no GFP peaks preprocessing (cf. section Microstate Measures). See also Fig. S2.1 for dynamic properties used in von Wegner et al., 2018.

As expected, the segmentation of the time series is very similar among three classical microstate algorithms (AAHC, TAAHC, and modified K-Means), and as seen in the previous section, this holds for both preprocessing options: with or without GFP peaks selection. These algorithms achieved accuracy scores of 0.9 (Fig. 8 left). On the other hand, the most dissimilar segmentation is between PCA and ICA, with an accuracy of around 0.2. The topographies (i.e., cluster centroids) are very similar across all six implemented algorithms, except for modified K-Means with GFP peaks preprocessing (Fig. 8 right). The three classical microstate algorithms produce similar static microstate statistics (coverage, lifespan, and occurrence) in both preprocessing options. The most dissimilar algorithm in the case of coverage is PCA and HMM. These two algorithms also produce dissimilar dynamic statistics, mainly entropy (Fig. S2.1).

### Subject-wise Synthetic Data

The next step in our investigation was to employ one step more realistic synthetic data, with more channels and mimicking the inter-subject variance, by generating 200 data samples, each with a different random seed for the random number generator. Therefore, each “subject” has a different positive-definite covariance matrix. To further extend our analysis and close the gap between the previous synthetic dataset and the following more realistic surrogate and, finally, the experimental dataset, we generated the following synthetic dataset with 20 channels and were seeking 4 microstates. Also, in this case, we only did use GFP peaks selection preprocessing, as is typical in microstate analyses. The overview of microstate properties for all six algorithms applied is presented in Fig. 9.

**Figure 9.**
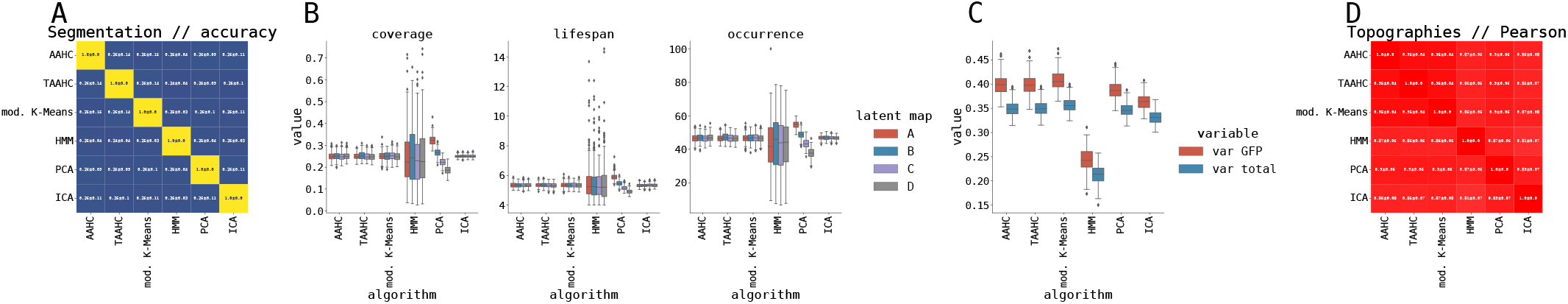
Microstate properties on the synthetic dataset with 200 simulated subjects using 6 algorithms for latent decomposition. (**A**) The similarity of segmentations as mean ± standard deviation across subjects as computed using 6 different microstate algorithms. The measure used to quantify similarity was accuracy. (**B**) Static microstate properties (coverage, lifespan, and occurrence; cf. section Microstate Measures) per latent map (A–D) and per algorithm. (**C**) Explained variance in GFP peaks and in entire time series per algorithm. (**D**) The similarity of topographies (cluster centres) yielded by each algorithm using Pearson correlation as mean ± standard deviation across subjects. See also Fig. S2.2 for dynamic properties used in von Wegner et al., 2018.

All tested algorithms are relatively stable with respect to modelling inter-subject variances (i.e., standard deviations on segmentation accuracy and topography correlations are low, and variances in box plots of different measures are also low) except for HMM. In particular, HMM shows relatively high inter-subject variance in coverage and occurrence (Fig. 9B), entropy, and entropy rate (Fig. S2.2). We also observe that 4 out of 6 algorithms on our synthetic data produce symmetric coverages (i.e., approximately 0.25 coverage for all 4 states; cf. Fig 9B), except for the aforementioned HMM and PCA (which orders components by their variance, hence it is natural that the coverages will be monotonically decreasing). A similar line of thought can be applied for lifespan and occurrences (Fig. 9B). We also note that the decomposition yielded by the three classical microstate algorithms (AAHC, TAAHC, and modified K-Means) explains the highest proportion of variance both in the GFP peaks and in the whole synthetic EEG time series, with PCA and ICA explaining slightly lower proportion of variance (Fig. 9C). The time series segmentation provided by all six algorithms exhibit similar mixing times, 4 out of 6 exhibit highest entropy (upper bound on the entropy is computed as the entropy of uniform distribution with 4 discrete states), with PCA being slightly lower, and HMM being significantly lower (this is connected with the coverages), see Fig. S2.2A. Finally, all six algorithms exhibit very similar first peaks of auto-mutual information (cf. Fig. S2.2A) and 5 out of 6 exhibit similar entropy rates (again, except for HMM; cf. Fig. S2.2B).

Summarizing the results from the synthetic dataset, we expect the three classical microstate algorithms to provide very similar topographies, time series segmentations, and thus both static and dynamic microstate properties.We also expect the HMM algorithm to perform differently than other tested algorithms, particularly in static microstate properties, the entropy of time series segmentation, and inter-subject variances in all measures. With these hypotheses in mind, let us move to the experiment dataset, i.e., resting-state eyes-open and eyes-closed EEG signal, acquired from 202 subjects.

### Experimental Data

In the experimental part of this study, we applied all 6 algorithms to the resting state EEG data we obtained for 202 subjects from the LEMON dataset (cf. section Experimental Data). All runs of our tested algorithms converged and provided a decomposition per subject and resting state paradigm (eyes-open and eyes-closed).

#### Microstate Maps

The so-called canonical microstate maps have been previously described for the AAHC and the modified K-Means algorithms (Dietrich Lehmann et al., 1987; Pascual-Marqui et al., 1995; Koenig et al., 2002). Koenig et al., 2002 provide these canonical maps as a binary data file, thus after each decomposition, we matched Koenig et al. microstate template maps with ours to make sure that our microstate A matches the template map A. Traditionally, the microstate A exhibits diagonal border between positive and negative potentials from the right frontal to the left occipital area. Microstate B exhibits a similar pattern, albeit the diagonal runs from the right frontal to the left occipital area. Microstate C has a horizontal orientation with a fronto-occipital gradient, while microstate D is often circular. We expect new geometries for some topographies to emerge for algorithms other than the classical ones (PCA, ICA, HMM). The overview of group topographies (i.e., the microstate maps) for the eyes-closed resting state EEG paradigm is shown in Fig. 10, while the eyes-open paradigm is shown in Fig. S1.1.

**Figure 10.**
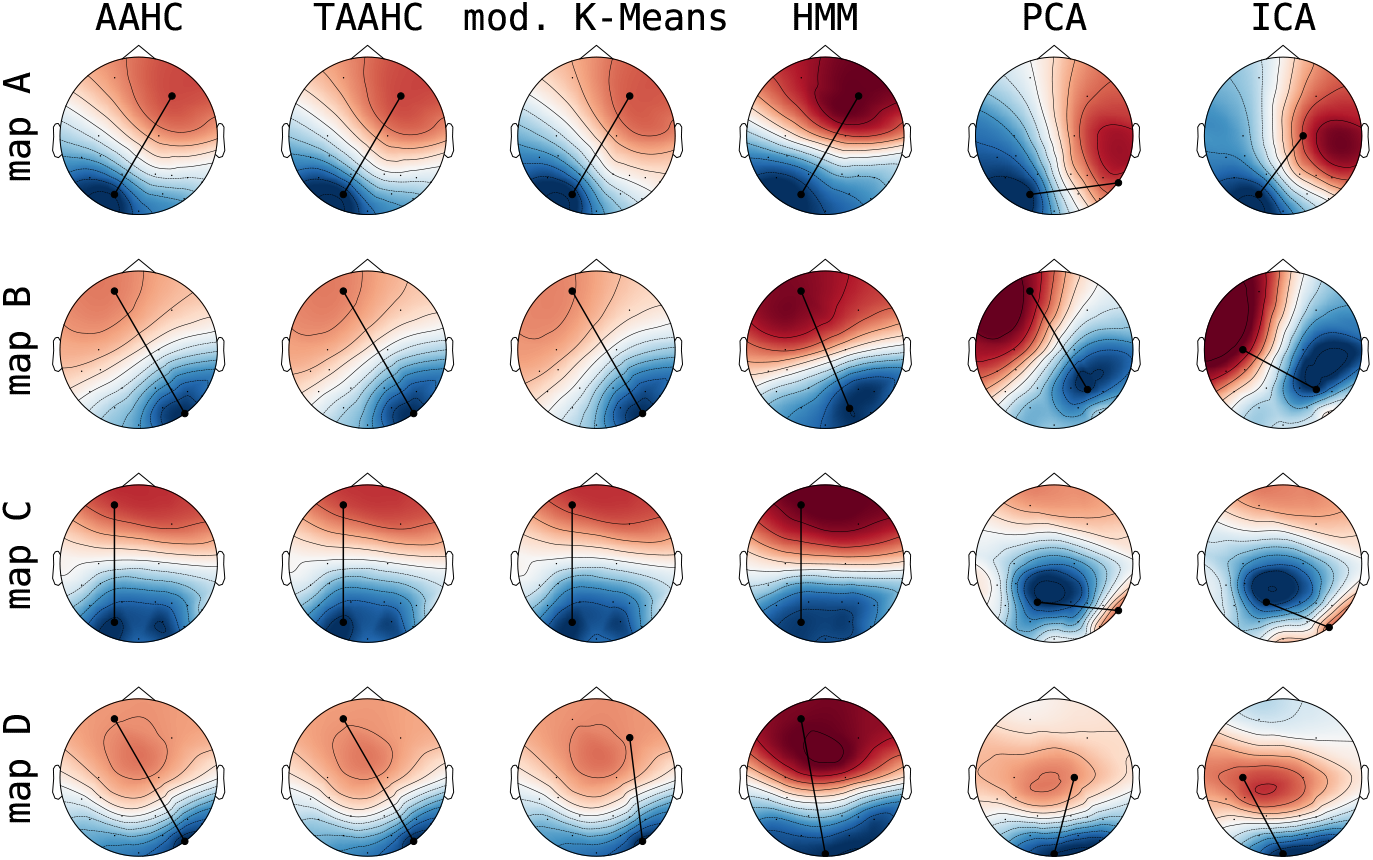
Group level microstate maps for LEMON dataset in the eyes-closed resting state EEG paradigm. The four microstates produced by each algorithm are shown row-wise, while different algorithms are shown column-wise. Group-level maps were computed by a simple average over all subjects with prior polarity matching based on microstate templates proved by Koenig et al., 2002. The maps also show a tangent between the maximum and minimum points on the map to guide readers to the orientation of the maximal gradient.

As expected, the classical microstate algorithms (AAHC, TAAHC, modified K-Means) show very similar geometries, and all bear a very close resemblance to the canonical microstate maps when all 4 states correlate with the templates at least at the 0.9 level. This is valid for both EEG paradigms (Fig. 10 and Fig. S1.1, first three columns), albeit the eyes-open paradigm exhibits a higher correlation on average. Other algorithms, namely the PCA, the ICA, and HMM show some subtle differences, albeit the main characteristics are conserved. In particular, all non-classical algorithms exhibit stronger gradients, mainly in maps A and B (cf. Fig. 10). Interestingly, most maps follow the classical description regarding the direction of gradients—this is true mainly for the HMM algorithm. Regarding PCA and ICA, maps A and B are somewhat similar to classical canonical microstates. However, maps C and D bear no resemblance. The same is valid for maps in the eyes-open case (Fig. S1.1).

Overall, we can conclude that the three classical microstate algorithms yield very similar topographies and correlate highly with the canonical template microstate maps (Koenig et al., 2002). The PCA and ICA topographies are somewhat similar, except for some multimodal maps and overall exhibit lower correlations with the canonical templates. Finally, the HMM-derived maps estimated via linear regression typically look very different, and most do not have a positive-negative potential gradient.

#### Microstate Properties

This section focuses on static and dynamic microstate properties resulting from 6 used algorithms. The overview plots are shown in Fig. 11 and Fig. S1.2 for the eyes-closed and eyes-open resting state EEG paradigm, respectively.

**Figure 11.**
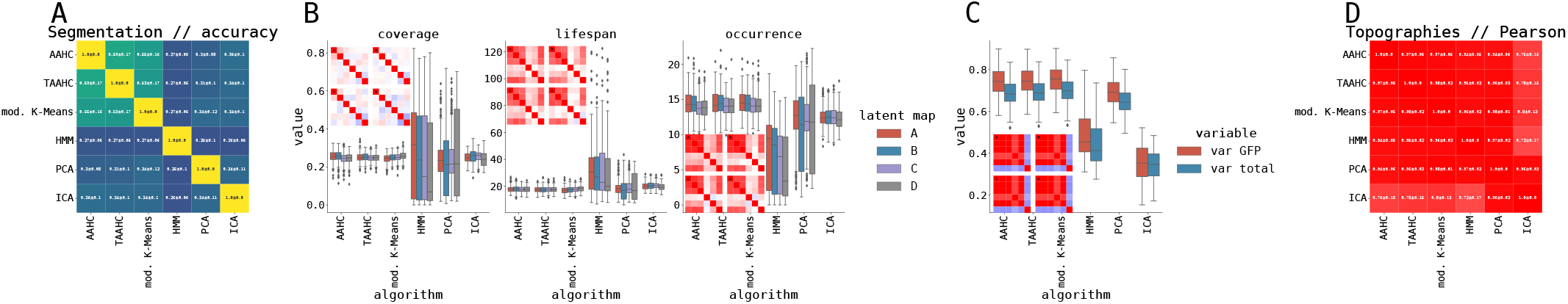
Overview of latent segmentation of a LEMON dataset in the eyes-closed resting state EEG paradigm. We computed a microstate segmentation, seeking 4 states in the experimental resting state EEG data in the eyes-closed paradigm (cf. section Experimental Data). In total, the dataset consists of 202 subjects. The first row shows: (**A**) The similarity of segmentations as mean ± standard deviation across subjects as computed using 6 different microstate algorithms. The measure used to quantify similarity was accuracy. (**B**) Static microstate properties (coverage, lifespan, and occurrence; cf. section Microstate Measures) per latent map (A–D) and per algorithm. Each plot contains a subplot, which visualises the correlation matrix of a given measure across subjects, averaged over A–D maps. (**C**) Explained variance in GFP peaks and in entire time series per algorithm. Shown is also the correlation matrix across subjects. (**D**) The similarity of topographies (cluster centres) yielded by each algorithm using Pearson correlation as mean ± standard deviation across subjects. See also Fig. S2.3 for dynamic properties used in von Wegner et al., 2018.

Compared with synthetic data, we expect more heterogeneity between individual subjects. This is immediately visible in the segmentation similarities (Fig. 11A): the three classical microstate algorithms provide the most similar segmentations with an average of accuracy (as a measure of similarity) between 0.5 and 0.6. Other algorithms show lower accuracy among themselves, as well as with regards to the three classical algorithms, averaging at approximately 0.3. The main question is whether this lower accuracy would create systematic differences in segmentation statistics such as coverage or lifespan. Fig. 11B shows that differences in segmentations are not particularly visible in statistics. In particular, the three classical microstate algorithms have virtually the same statistics (coverage, lifespan, and occurrence), and the correlation across subjects is very high. In other algorithms, we see some systematic differences. HMM and PCA exhibit high variance across subjects in all three statistics, and the PCA algorithm shows a low or even negative correlation with respect to other tested algorithms. Finally, ICA is more or less similar to the three classical algorithms, with lower correlations across subjects. The decomposition done by the three classical algorithms also explains the highest proportion of variance in the original EEG (Fig. 11C). Final topographies (i.e., microstates or cluster centroids) are largely similar between algorithms (approximately 0.9) with the little exception of ICA (0.7), cf. Fig. 11D. Regarding dynamic properties (Fig. S2.3A), all others exhibit similar mixing times except for the HMM algorithm. PCA and HMM algorithms also show lower entropy and entropy rate values, while all algorithms agree on the first peak of the auto-mutual information function.

In summary, the three classical microstate algorithms largely provide the same results of static and dynamic microstate statistics and topographies. The differences in entire time series segmentations (accuracy approximately 0.5–0.6) are smeared out and do not propagate to any other statistics. The most dissimilar algorithm is the hidden Markov Model, which exhibits much higher subject-wise variance in static microstate properties, lower entropy and entropy rate values, and the lowest overall accuracy (with respect to other algorithms) of time series segmentation. PCA and ICA share some properties with HMM (large subject-wise variance in static microstate properties, lower explained variance).

#### Comparison with Synthetic Data

Here we shortly compare the results from our synthetic data pipeline with microstate analysis on real resting-state EEG data (i.e., Fig. 9 and Fig. 11). The analysis of our synthetic datasets partially explains the differences between algorithms. The three classical microstate algorithms exhibit very similar statistics in both cases. HMM, in both cases, possesses significantly higher subject-wise variance in static microstate statistics. Explained variance is the highest for the three classical algorithms and the lowest for HMM. HMM and PCA segmentations exhibit lower entropy and entropy rates. However, the similarity of topographies differs between results for EEG data and synthetic data: for resting-state EEG, the topographies are, overall, more similar between the methods, and we can also observe the division between the three classical microstate algorithms and the three other. This division is not visible in our synthetic data, and all off-diagonal elements in the accuracy matrix have approximately the same, relatively low, value.

### Surrogate Data

The next in our investigation was to run the microstate analysis on surrogate data. We compute multivariate Fourier Transform (FT) surrogate data (Theiler et al., 1992) from resting-state EEG for each subject separately. FT surrogate is a randomization procedure where the original data is transformed into the Fourier spectral domain, its phases are randomized, and then the data with random Fourier phases are transformed back into the time domain. Moreover, the randomization is the same across channels as we use a multivariate version of the FT procedure. This ensures that the surrogate data possess the same covariance and temporal autocorrelation structure as the original data, but any (nonlinear) dependencies of higher order are destroyed. Should the results from microstate analysis on FT surrogate data be comparable to the original resting-state EEG, we conclude that the microstate properties are well explained by the covariance and autocorrelation structure of the data and, as such, might be estimated from these.

The group-level topographies from microstate analysis on FT surrogate data are similar to the resting-state EEG data (cf. Fig S1.3)since the covariance structure and phase mainly determine their randomization preserves covariances. However, the microstate statistics could still differ, as shown in Fig. 12.

**Figure 12.**
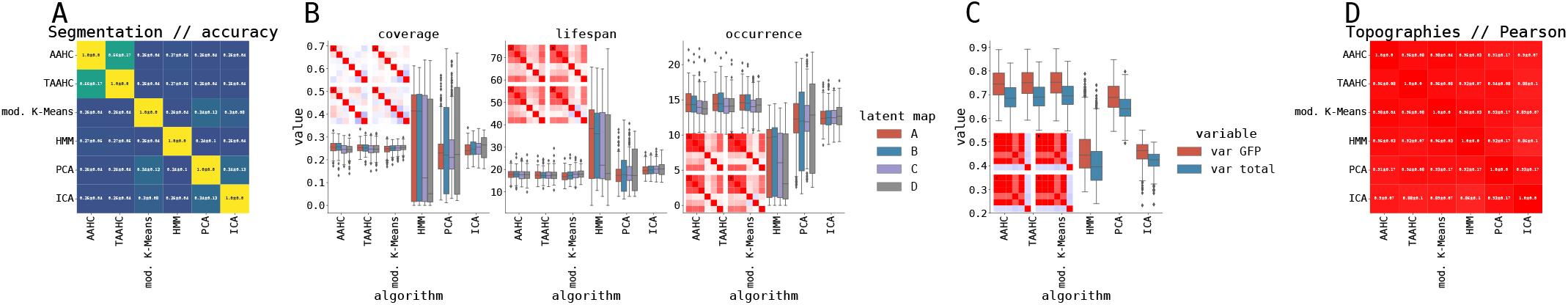
Overview of latent segmentation of a Fourier Transform surrogate data generated from the LEMON dataset in the eyes-closed resting state EEG paradigm. We computed a microstate segmentation, seeking 4 states in the experimental resting state EEG data in the eyes-closed paradigm (cf. section Experimental Data). In total, the dataset consists of 202 subjects. The first row shows: (**A**) The similarity of segmentations as mean ± standard deviation across subjects as computed using 6 different microstate algorithms. The measure used to quantify similarity was accuracy. (**B**) Static microstate properties (coverage, lifespan, and occurrence; cf. section Microstate Measures) per latent map (A–D) and algorithm. Each plot contains a subplot, which visualises the correlation matrix of a given measure across subjects, averaged over A–D maps. (**C**) Explained variance in GFP peaks and in entire time series per algorithm. Shown is also the correlation matrix across subjects. (**D**) The similarity of topographies (cluster centres) yielded by each algorithm using Pearson correlation as mean ± standard deviation across subjects. See also Fig. S2.4 for dynamic properties used in von Wegner et al., 2018.

The segmentation accuracy exhibits similar values as in the real EEG data case. However, we note that for the FT surrogates, only the AAHC and TAAHC exhibit mutually higher accuracies, while modified K-Means have similar accuracy as other algorithms (Fig. 11A vs Fig. 12A). However, the static and widely used microstates statistics such as coverage, lifespan, and occurrence are very well reproduced by FT surrogates with the same patterns: three classical microstate algorithms in a general agreement and with low subject-wise variance, and HMM and PCA having large subject-wise variance (Fig. 11B and Fig. 12B). Even the correlation across subjects show the same patterns.

Similarly, the explained variance by the microstate segmentation exhibits the same patterns, with the three classical algorithms explaining the highest portion of the variance, followed by the PCA and finally, ICA and HMM (Fig. 11C and Fig. 12C). Virtually the same results are also obtained from the dynamic statistics such as mixing time, entropy, entropy rate, and the first peak of auto-mutual information function (cf. Fig. S2.3A and B and Fig. S2.4A and B).

In summary, except for minor deviations for the modified K-Means when looking at the mutual accuracy of the segmentations, all other microstate statistics (both static and dynamic) seem to be very well reproduced by the FT surrogates of the real resting-state EEG data. This indicates that covariance structure and temporal autocorrelation may substantially contribute to these observed microstate statistics. The actual computationally demanding microstate decomposition might provide only limited added value with respect to estimating these quantities or further using them to characterise the brain connectivity dynamics.

### Vector Autoregression Data

Our experiment with swapping real resting-state EEG data with its FT surrogate yielded virtually the same results, leading us to our final hypothesis and experiment. Suppose the microstate statistics are mainly given by the covariance and autocorrelation structure. In that case, fitting a Vector Autoregression (VAR) process (cf. section Vector Autoregression Data) and generating a simulation with additive Gaussian noise should yield the same microstate statistics. To this end, we designed the following experiment. First, for each subject, we fitted a VAR process of order *p* with *p* ∈ [0, …, 20] and found the order that minimizes the Akaike information criterion. Then we took a median over optimized orders *p* to determine the overall best order for all subjects (for *p* > 5, AIC changed only very slightly, and we deemed the overall best order as 8, not shown). Secondly, we chose a length of the data segment, *t*_*seg*_ ∈ [10, 30, 60, 180] seconds and for all subjects, we computed microstate decomposition and statistics using modified K-Means algorithm for two consecutive resting-state EEG data segments (each of length *t*_*seg*_), for simulated VAR process of total length *t*_*V AR*_ = 3600 seconds, and for all individual segments of length *t*_*seg*_ of the long VAR simulation (e.g., for *t*_*seg*_ = 10 we had 360 segments).

Having computed microstate decompositions and statistics for these 4 cases (two consecutive EEG data segments, long VAR process, and individual segments of long VAR process), we evaluated the “prediction error” (i.e., the difference in microstate statistics estimation measured by mean square error, MSE) between the first and the second data segment, between the mean of VAR segments and the second data segment, and between the long VAR process and the second data segment. Finally, we compared these prediction errors (MSE in the estimation of microstate properties) between these three cases using pairwise paired t-tests, i.e. evaluating contrasts: data prediction error vs VAR segments prediction error, data prediction error vs full VAR prediction error, and full VAR prediction error vs VAR segments prediction error. Fig. 13 shows T-values and p-values for the contrast between VAR segments prediction error and data prediction error, while Fig. S1.4 shows T-values and p-values for the other two contrasts. Both figures show these metrics for all four tested segment lengths.

**Figure 13.**
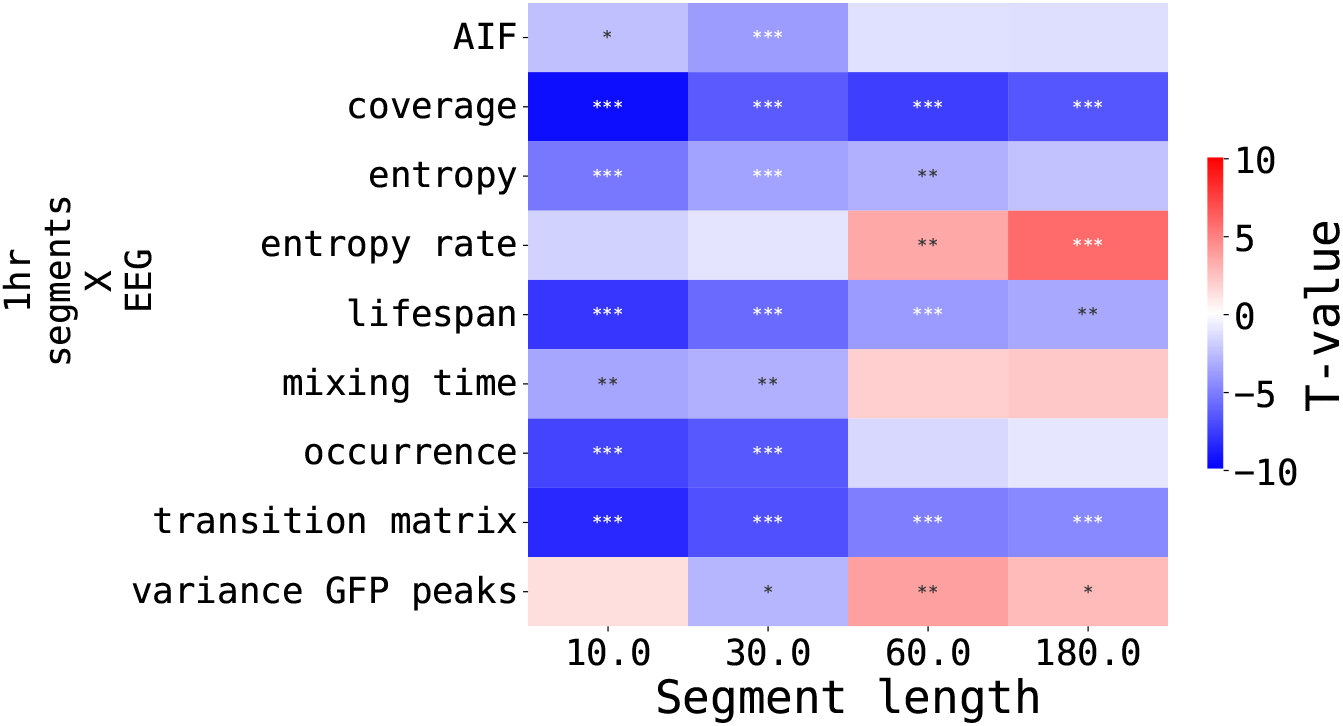
VAR segments microstate estimation comparison with real EEG data as t-tests on a mean square error in short segments. T-tests were computed on a sample of “prediction errors” between the first and the second data segment per subject and the average of “prediction errors” between segments of the VAR process and the second data segment per subject (see main text for details). The contrast was computed as VAR segments vs real data, i.e., negative T-values (blue) indicate lower MSE for the VAR process. The plot colour-codes the T-value and stars encode statistical significance (* for p-values < 0.05, ** for p-values < 0.01 and * * * for p-values < 0.001).

As Fig. 13 clearly shows, most measures (except entropy rate and explained variance) were better “predicted” from VAR segments. By predicted, we mean that the VAR process can generate dynamics closer to the new data segment than the previous data segment, at least in the view of microstate statistics. This holds for all lengths of the segments. Naturally, for very short segments (10 seconds), the overall stability of the microstate estimate would not be particularly high. Entropy rate and mixing time measures were well estimated by VAR segments for short segment lengths (10 and 30 seconds). However, for longer segments (60 and 180 seconds), the first data segment was closer to the second data segment than the VAR segments: for longer segment lengths, the estimate of entropy rate and mixing time is better and converges to its theoretical value. Overall, all static and some dynamic microstate properties were correctly estimated using the population average of VAR segments, which points to the fact that microstate properties and statistics are indeed determined mainly by the covariance and temporal autocorrelation structure.

## Discussion

In the first part of our study, we thoroughly investigated 6 microstate algorithms, firstly by estimating microstate properties on synthetic Gaussian data (cf. section Understanding Clustering Algorithms on Synthetic Data) and then by estimating topographies and microstate properties of real EEG data (cf. section Experimental Data). A similar endeavour was already partially undertaken by von Wegner et al., 2018. However, we added the HMM algorithm as an option for clustering data into microstates and added experiments with synthetic data in order to understand the dynamics of the clustering. Moreover, we also utilised FT surrogates and a VAR model that underwent the same analysis in order to test our hypothesis that microstate properties are primarily dependent on the linear structure of the underlying EEG data.

Our first observation is a minimal effect of temporal reduction of the data by computing GFP peaks. The only exception is the HMM algorithm (see Fig. 5). Importantly, we do recommend using GFP peaks subsampling, particularly for classical bottom-up approaches to microstate clustering, i.e., AAHC and TAAHC algorithms, since they are computationally very demanding without an option for straightforward parallelisation and thus cannot exploit advances in affordable multi-CPU setups.

We are aware of the main limitation of our approach for trying to visualise and understand geometrically the workings of and similarities of various microstate algorithms with synthetic data (cf. section Synthetic Data), and that is that real EEG data are not purely Gaussian. However, such a simple model allows for geometric insights and provides predictions that, in many cases, were reflected in real data. Moreover, we tried various dimensionality reduction schemes (ranging from linear PCA to nonlinear t-SNE (Hinton et al., 2002) and locally linear embedding (Roweis et al., 2000)) and observed the distribution of embeddings that are very close to the multivariate normal distribution in the first 5 reduced dimensions (not shown). Therefore, we conclude that this approximation is relatively accurate, and the approximation cost is negligible. Finally, the inherent notion of Gaussianity is also present in our FT surrogate and, ultimately, VAR model results (cf. sections Vector Autoregression Data and Vector Autoregression Data), where we observed, that indeed, such an approximate linear/Gaussian model is able to capture microstate properties of real EEG data to a surprisingly large extent.

Our hypothesis of microstate properties being determined mainly by the underlying data’s linear structure is somewhat shown by our experiments with FT surrogates (cf. section Surrogate Data) and VAR process (cf. section Vector Autoregression Data). We note that we do not provide any analytical treatment that would rigorously prove our hypothesis, and this is beyond the scope of our current study, which aims to be purely computational. These computational experiments strongly hint at the fact that static and dynamic microstate properties depend to a very high degree on the linear covariance and autocorrelation structure of the underlying EEG data. Multivariate FT surrogates strictly preserve both autocorrelation of EEG channels and their covariance structure, and the estimated microstate properties were practically identical to those of real data (compare Figs. 11 and 12).

Our final experiment with estimating microstate properties from the VAR process proved that most measures (except entropy rate and explained variance) are comparable to those estimated from real EEG data (cf. section Vector Autoregression Data). There were two exceptions, though: mixing time and entropy rate because the estimation of these variables largely depends on the length of the simulated VAR segment. To summarise, all static and some dynamic microstate properties were correctly estimated using the population average of VAR segments, which points to the fact that microstate properties and statistics are indeed determined mainly by the covariance and temporal autocorrelation structure.

Finally, we conclude with a few critical points about the methodology of how microstate analysis is used throughout the literature. We discovered that the preprocessing using GFP peaks does not alter either microstate segmentation or resulting topographies, i.e., cluster centres. However, we note that it is beneficial for classical bottom-up algorithms AAHC and TAAHC purely from the perspective of the computational burden. On the other hand, we observed that all microstate algorithms found 2 clusters in our synthetic data example (cf. section Understanding Clustering Algorithms on Synthetic Data) even when the data were coming from a multivariate Gaussian distribution. There was an exception with the HMM algorithm: without the preprocessing with GFP peaks, it correctly attributed all temporal points to one cluster while the second was empty. However, after the GFP peaks finding procedure (which selects data points further away from the mean vector), HMM could also find two clusters in the data. In other words, the GFP finding procedure introduced an artificial manifold separating the data from the same distribution. Lastly, we question the necessity and advantages of microstate analysis: some microstate properties are tied to clinical findings. However, if these properties are determined by the covariance and temporal autocorrelation structure of the data, we should, in principle, be able to tie these clinical findings to linear properties of the EEG data (such as the spectral power or covariances) without the need to undergo the computationally more demanding and likely less robust microstate analysis.

## Conclusion

The main aim of this study was to dive deep into the exact dynamics of microstate analysis to systematically compare various algorithms typically used in the literature. Except for HMM, all other 5 selected algorithms arrived at similar results and did not show significant differences.

In the next step, we systematically compared clustering algorithms by running a complete microstate analysis on publicly available EEG data and computed typically used static and dynamic statistics. In line with previous works in this area, we conclude no significant differences between the algorithms except for HMM.

Finally, we tested our hypothesis that most microstate properties are determined purely by the covariance and autocorrelation structure of the underlying EEG signal; firstly, by generating a multivariate Fourier Transform surrogate of EEG data and performing microstate analysis, and secondly, by fitting a Vector Autoregression process to the EEG data, generating a sample from the fitted VAR process, and estimating microstate properties on VAR segments of various lengths. We observed that most measures (except entropy rate and explained variance) were better “predicted” from VAR segments. By predicted, we mean that the VAR process can generate dynamics closer to the data segment than the previous data segment, at least in the view of microstate statistics.

## Acknowledgment

This preprint was created using the LaPreprint template (https://github.com/roaldarbol/lapreprint) by Mikkel Roald-Arbøl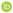. Computational resources were supplied by the project “e-Infrastruktura CZ” (e-INFRA CZ LM2018140) supported by the Ministry of Education, Youth and Sports of the Czech Republic. This project has received funding from the European Union’s Horizon Europe research and innovation programme under the HORIZON-WIDERA grant agreement No. 101090306 (N.J.) and from the Czech Science Foundation project No. 21-32608S (J.H.).

## Author contributions

**N.J**.: conceptualisation, methodology, code, formal analysis, visualisation, writing, editing. **J.H**.: conceptualisation, methodology, resources, writing, editing, supervision.

## Appendix 1

### Supplementary figures

#### Eyes open resting state paradigm in real data

**Appendix 1—figure S1.1.**
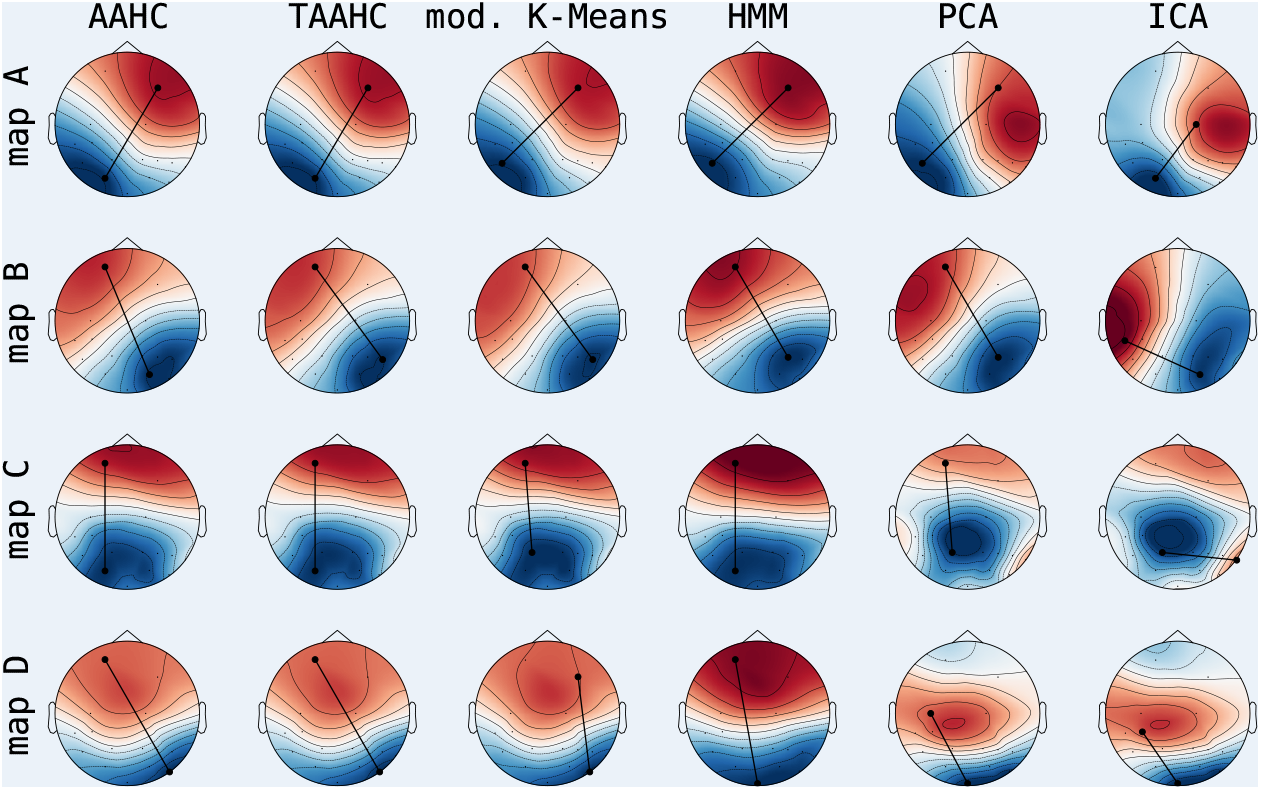
Group-level microstate maps for LEMON dataset in the eyes-open resting state EEG paradigm. The four microstates produced by each algorithm are shown row-wise, while different algorithms are shown column-wise. Group-level maps were computed by a simple average over all subjects with prior polarity matching based on microstate templates proved by Koenig et al., 2002. The maps also show a tangent between the maximum and minimum points on the map to guide readers to the orientation of the maximal gradient.

**Appendix 1—figure S1.2.**
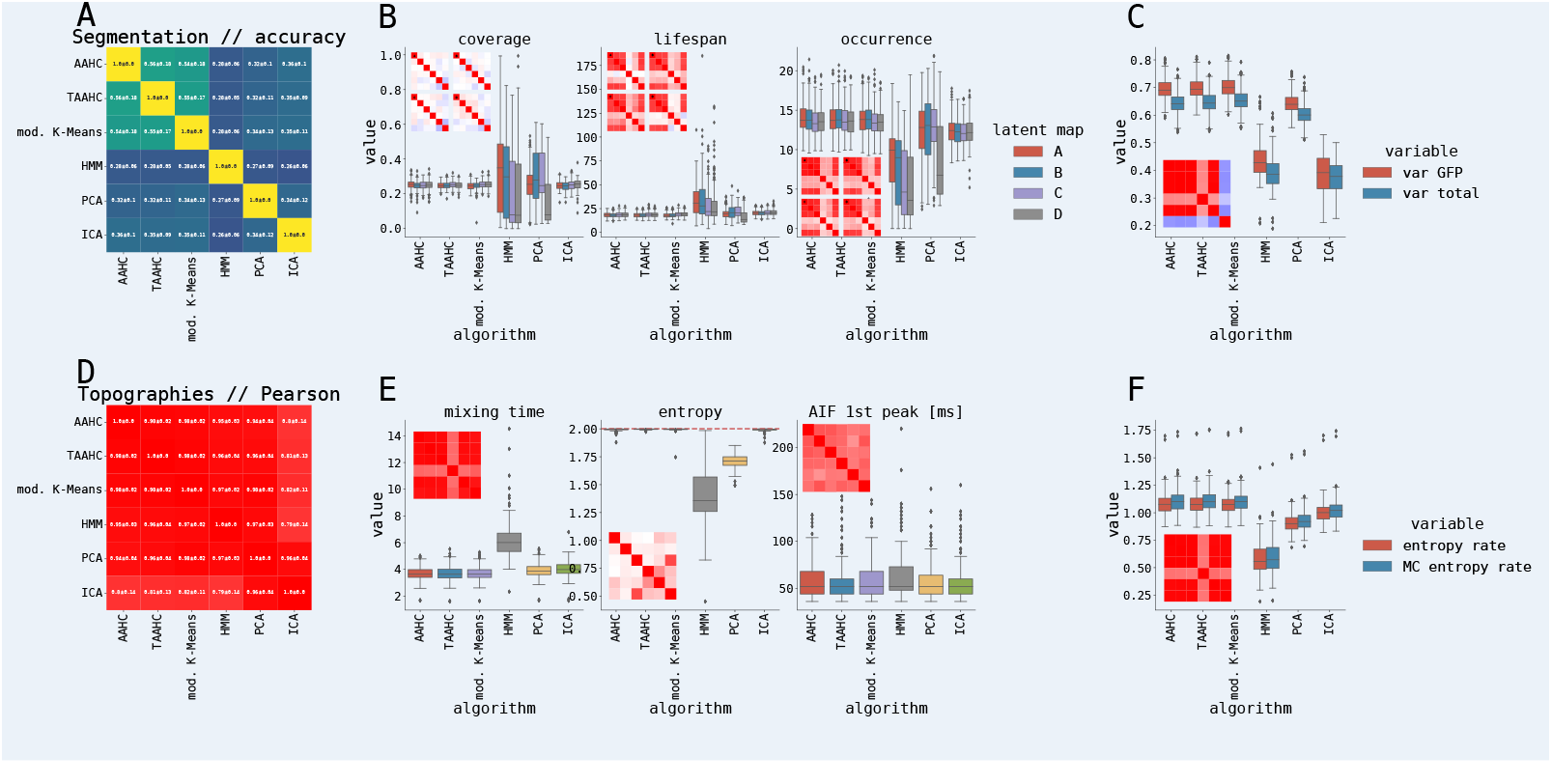
Overview of latent segmentation of a LEMON dataset in the eyes-open resting state EEG paradigm. We computed a microstate segmentation, seeking 4 states in the experimental resting state EEG data in the eyes-open paradigm (cf. section Experimental Data). In total, the dataset consists of 202 subjects. The first row shows: (**A**) The similarity of segmentations as mean ± standard deviation across subjects as computed using 6 different microstate algorithms. The measure used to quantify similarity was accuracy. (**B**) Static microstate properties (coverage, lifespan, and occurrence; cf. section Microstate Measures) per latent map (A–D) and algorithm. Each plot contains a subplot, which visualises the correlation matrix of a given measure across subjects, averaged over A–D maps. (**C**) Explained variance in GFP peaks and in entire time series per algorithm. Shown is also the correlation matrix across subjects. (**D**) The similarity of topographies (cluster centres) yielded by each algorithm using Pearson correlation as mean ± standard deviation across subjects. (**E**) dynamic statistics (cf. section Information-Theoretical Analysis) of the segmentations: mixing time, entropy (red dashed line shows maximum entropy given the number of states) and a first peak of auto-information function, and (**F**) the entropy rate of an actual decomposition (red) and the entropy rate of idealised Markov process given the empirical transition matrix (blue).

#### Surrogate Data Microstate Maps

**Appendix 1—figure S1.3.**
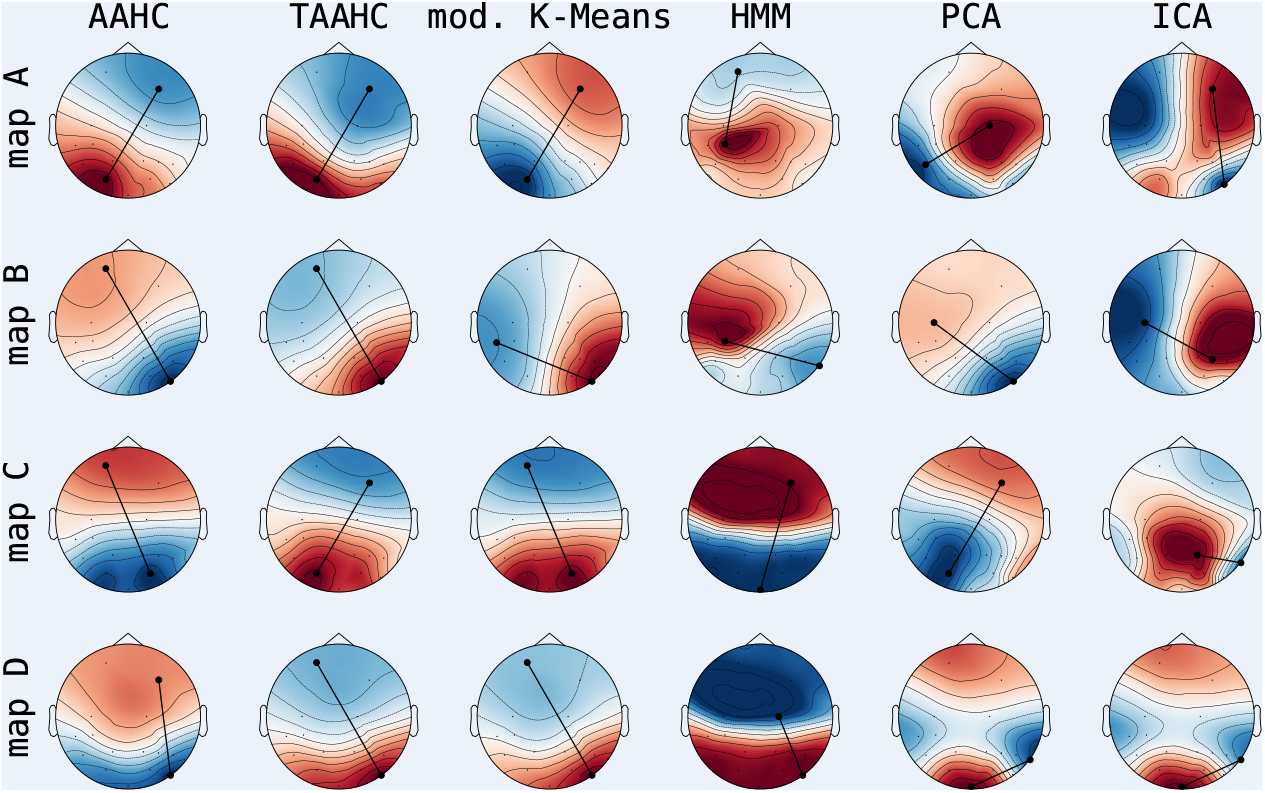
Group-level microstate maps for Fourier Transform surrogates generated from the LEMON dataset in the eyes-closed resting state EEG paradigm. The four microstates produced by each algorithm are shown row-wise, while different algorithms are shown column-wise. Group-level maps were computed by a simple average over all subjects with prior polarity matching based on microstate templates proved by Koenig et al., 2002. The maps also show a tangent between the maximum and minimum points on the map to guide readers to the orientation of the maximal gradient.

#### Full length VAR process comparison with EEG data and VAR process segments

**Appendix 1—figure S1.4.**
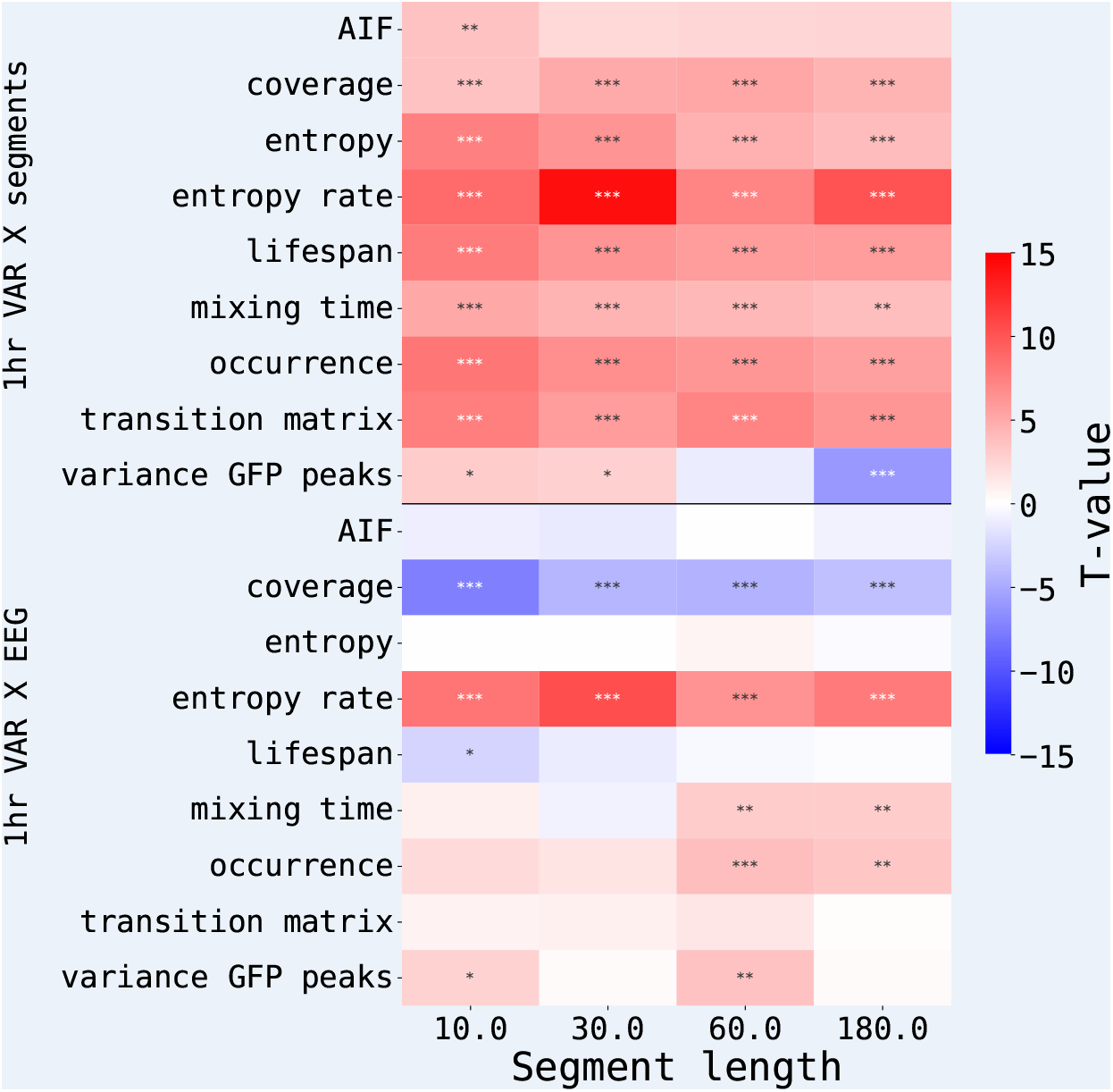
VAR process microstate estimation comparison with real EEG data and with VAR segments as t-tests on a mean square error in short segments. T-tests were computed on a sample of “prediction errors” between the first and the second data segment per subject, the average of “prediction errors” between segments of the VAR process and the second data segment per subject, or between full VAR process and second data segment per subject (see main text for details). The contrasts were computed as full VAR vs VAR segments and vs real data, i.e., negative T-values (blue) indicate lower MSE for the full VAR process. The plot colour-codes the T-value and stars encode statistical significance (* for p-values < 0.05, ** for p-values < 0.01 and * * * for p-values < 0.001).

## Appendix 2

### Information-Theoretical Analysis

#### Methods

In addition to classical microstate measures presented in the main text, we also computed a number of statistics stemming from the information-theoretical analysis, following Wegner et al., 2017 and von Wegner et al., 2018. Here we again operate on the symbolic state sequences *u*_*t*_, resulting from clustering algorithms. In the following text, we denote *p* the empirical distribution of microstate labels (i.e., the distribution of coverages, cf. eq. (5)), and *T* the empirical transition matrix between states.

##### Entropy

Shannon entropy characterises information content in a sequence (Kullback, 1959), with *h* = 0 being its minimum value meaning no randomness and a delta distribution. On the other hand, maximum Shannon entropy *h* = log *m* with *m* being a number of possible states describes a uniform distribution with maximum randomness. Shannon entropy is computed as

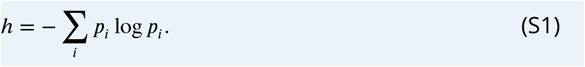

The units of entropy are bits or nats, depending on the base of logarithms (binary and natural, respectively).

##### Entropy rate

The entropy rate 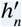 of a stochastic process quantifies how much uncertainty the process produces at each new time step, given the information about the past states (Levin et al., 2006) (valid in the limit of *n* → ∞):

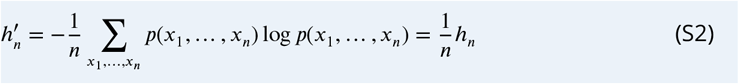

where *p*(*x*_1_, …, *x*_*n*_) denotes the joint probability of a specific sequence (*x*_1_, …, *x*_*n*_). To approximate eq. (S2), the entropy rate 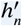 is estimated as the slope of the linear least squares fit of *n* vs *h*_*n*_ (Lizier et al., 2012). In our estimations, we reused the history parameter *n* = 6 following von Wegner et al., 2018.

##### Mixing time

The mixing time *τ* of a Markov process summarises a transition matrix of a process in a single number. Mixing time is closely related to a spectral gap computed as a difference between the largest and second largest eigenvalues of a transition matrix, *λ*_0_ − *λ*_1_. Since *λ*_0_ = 1 for stochastic matrices as per the Perron-Frobenius theorem (von Wegner et al., 2018), the mixing time is thus defined as

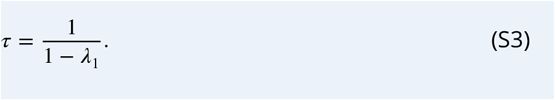

##### Auto-information peaks

The dominant periodicity of a microstate sequence is summarised by the location of the first nontrivial peak of the auto-information function *I*(*k*), where

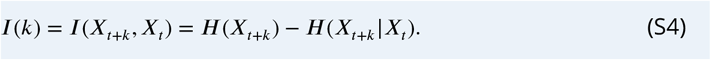

Auto-information function always has the global maximum at the time lag 0; hence we seek local maxima after the minimum of 8 time steps (i.e., 32 ms with a sampling rate of 250 Hz).

To reduce the influence of noise, we smooth the auto-information function with a moving average filter of size 3 time steps, in line with von Wegner et al., 2018.

#### Synthetic Data Microstate Properties

**Appendix 2—figure S2.1.**
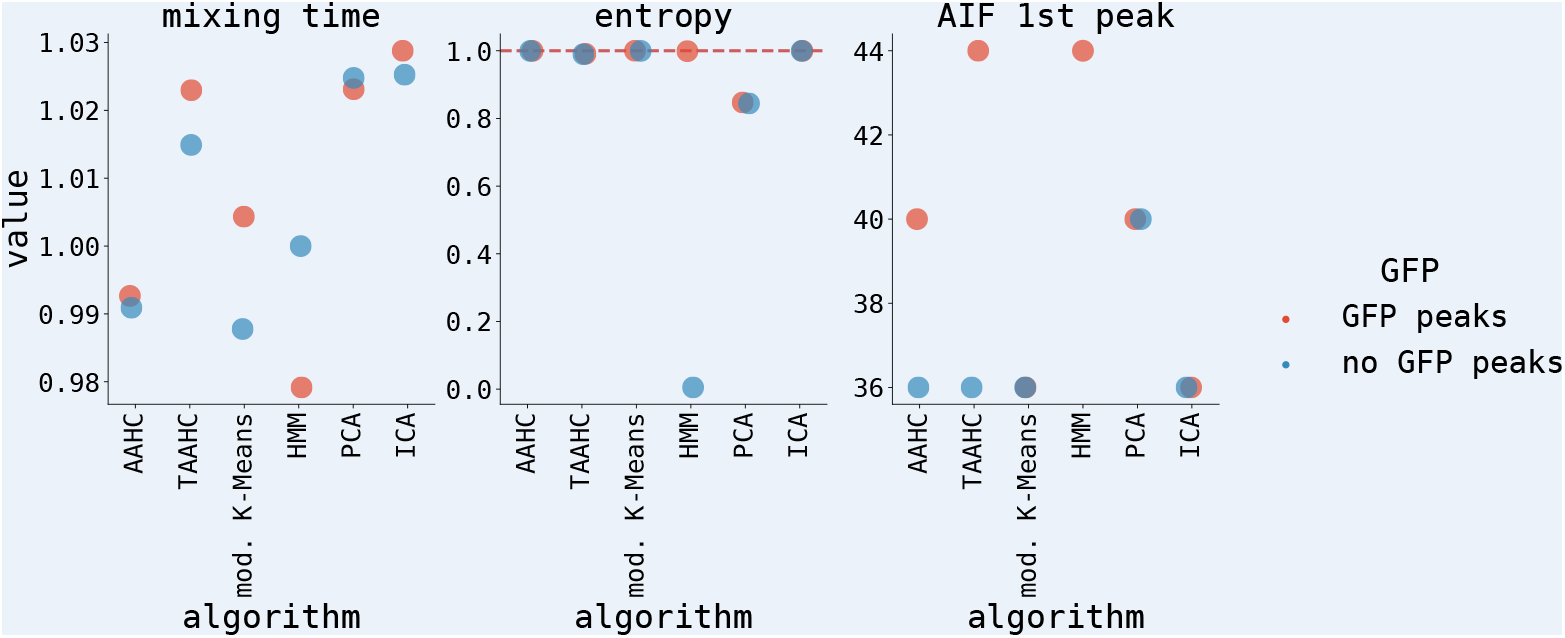
Microstate properties on the synthetic dataset using 6 algorithms for latent decomposition. The figure shows mixing time, entropy and the first peak of the auto-mutual information function (cf. section Information-Theoretical Analysis)

#### Subject-wise Synthetic Data

**Appendix 2—figure S2.2.**
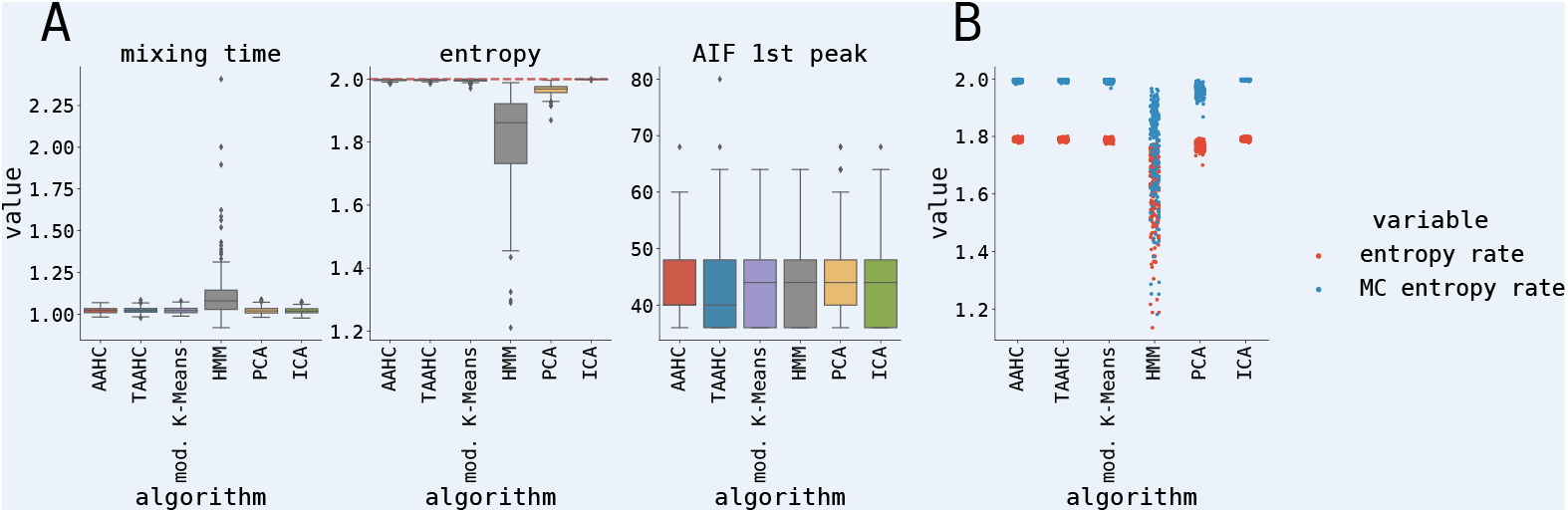
Microstate properties on the synthetic dataset with 200 simulated subjects using 6 algorithms for latent decomposition. (**A**) Dynamic microstate properties (mixing time, entropy, and the first peak of the auto-mutual information function; cf. section Information-Theoretical Analysis) per algorithm. (**B**) Entropy rate and idealized entropy rate of a Markov Chain process with the same empirical transition matrix as synthetic data per algorithm.

#### Experimental Data: Microstate Properties

**Appendix 2—figure S2.3.**
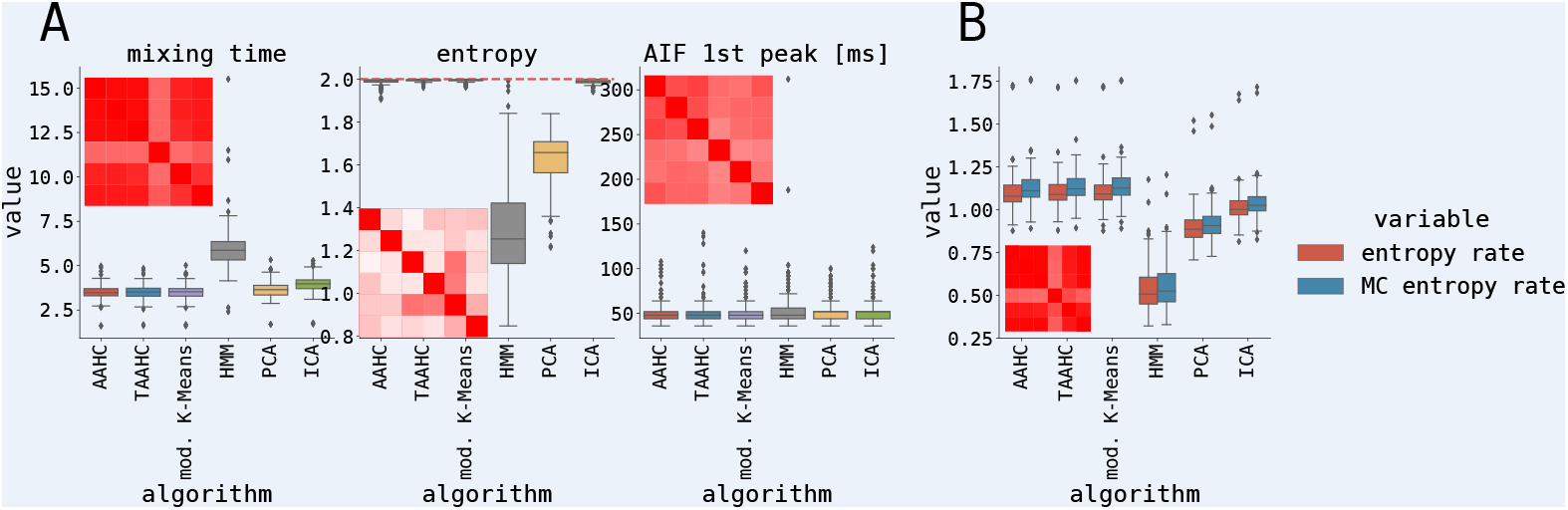
Overview of latent segmentation of a LEMON dataset in the eyes-closed resting state EEG paradigm. (**A**) Dynamic microstate properties (mixing time, entropy, and the first peak of the auto-mutual information function; cf. section Information-Theoretical Analysis) per algorithm. Shown are also correlation matrices of given measures across subjects. (**B**) Entropy rate and idealized entropy rate of a Markov Chain process with the same empirical transition matrix as synthetic data per algorithm.

#### Surrogate Data

**Appendix 2—figure S2.4.**
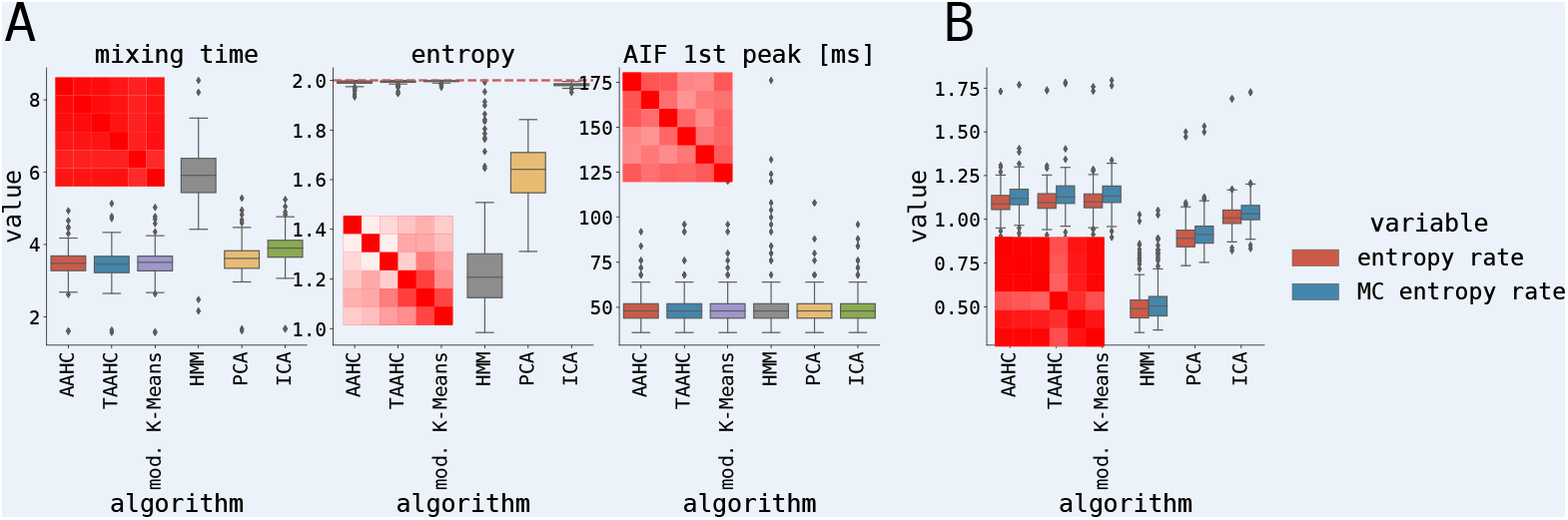
Overview of latent segmentation of a Fourier Transform surrogate data generated from the LEMON dataset in the eyes-closed resting state EEG paradigm. (**A**) Dynamic microstate properties (mixing time, entropy, and the first peak of the auto-mutual information function; cf. section Information-Theoretical Analysis) per algorithm. Shown are also correlation matrices of given measures across subjects. (**B**) Entropy rate and idealized entropy rate of a Markov Chain process with the same empirical transition matrix as synthetic data per algorithm.

## References

Babayan, Anahit, Miray Erbey, Deniz Kumral, Janis D Reinelt, Andrea MF Reiter Josefin Röbbig, H Lina Schaare, Marie Uhlig, Alfred Anwander, Pierre-Louis Bazin, et al. (2019). “A mind-brain-body dataset of MRI, EEG, cognition, emotion, and peripheral physiology in young and old adults”. In: Scientific data 6.1, pp. 1–21.

Baker, Adam P, Matthew J Brookes, Iead A Rezek, Stephen M Smith, Timothy Behrens, Penny J Probert Smith, and Mark Woolrich (2014). “Fast transient networks in spontaneous human brain activity”. In: elife 3, e01867.

Beckmann, Christian F, Marilena DeLuca, Joseph T Devlin, and Stephen M Smith (2005). “Investigations into resting-state connectivity using independent component analysis”. In: Philosophical Transactions of the Royal Society B: Biological Sciences 360.1457, pp. 1001–1013.

Britz, Juliane, Dimitri Van De Ville, and Christoph M Michel (2010). “BOLD correlates of EEG topography reveal rapid resting-state network dynamics”. In: Neuroimage 52.4, pp. 1162–1170.

Chaudhuri, Rishidev, Berk Gerçek, Biraj Pandey, Adrien Peyrache, and Ila Fiete (2019). “The intrinsic attractor manifold and population dynamics of a canonical cognitive circuit across waking and sleep”. In: Nature neuroscience 22.9, pp. 1512–1520.

Deco, Gustavo and Morten L Kringelbach (2016). “Metastability and coherence: extending the communication through coherence hypothesis using a whole-brain computational perspective”. In: Trends in neurosciences 39.3, pp. 125–135.

Fox, Michael D and Marcus E Raichle (2007). “Spontaneous fluctuations in brain activity observed with functional magnetic resonance imaging”. In: Nature reviews neuroscience 8.9, pp. 700–711.

Gallego, Juan A, Matthew G Perich, Lee E Miller, and Sara A Solla (2017). “Neural manifolds for the control of movement”. In: Neuron 94.5, pp. 978–984.

Gallego, Juan A, Matthew G Perich, Stephanie N Naufel, Christian Ethier, Sara A Solla, and Lee E Miller (2018). “Cortical population activity within a preserved neural manifold underlies multiple motor behaviors”. In: Nature communications 9.1, p. 4233.

Gramfort, Alexandre, Martin Luessi, Eric Larson, Denis A. Engemann, Daniel Strohmeier, Christian Brodbeck, Roman Goj, Mainak Jas, Teon Brooks, Lauri Parkkonen, and Matti S. Hämäläinen (2013). “MEG and EEG Data Analysis with MNE-Python”. In: Frontiers in Neuroscience 7.267, pp. 1–13.

Haykin, Simon and N Network (2004). “A comprehensive foundation”. In: Neural networks 2.2004, p. 41.

Hinton, Geoffrey E and Sam Roweis (2002). “Stochastic neighbor embedding”. In: Advances in neural information processing systems 15.

Hyvärinen, Aapo and Erkki Oja (2000). “Independent component analysis: algorithms and applications”. In: Neural networks 13.4-5, pp. 411–430.

Jazayeri, Mehrdad and Arash Afraz (2017). “Navigating the neural space in search of the neural code”. In: Neuron 93.5, pp. 1003–1014.

Khanna, Arjun, Alvaro Pascual-Leone, and Faranak Farzan (2014). “Reliability of resting-state microstate features in electroencephalography”. In: PloS one 9.12.

Kikuchi, Mitsuru, Thomas Koenig, Yuji Wada, Masato Higashima, Yoshifumi Koshino, Werner Strik, and Thomas Dierks (2007). “Native EEG and treatment effects in neuroleptic-naive schizophrenic patients: time and frequency domain approaches”. In: Schizophrenia research 97.1-3, pp. 163–172.

Kiviniemi, Vesa, Juha-Heikki Kantola, Jukka Jauhiainen Aapo Hyvärinen, and Osmo Tervonen (2003). “Independent component analysis of nondeterministic fMRI signal sources”. In: Neuroimage 19.2, pp. 253–260.

Koenig, Thomas, Dietrich Lehmann, Marco CG Merlo, Kieko Kochi, Daniel Hell, and Martha Koukkou (1999). “A deviant EEG brain microstate in acute, neuroleptic-naive schizophrenics at rest”. In: European archives of psychiatry and clinical neuroscience 249.4, pp. 205–211.

Koenig, Thomas, Leslie Prichep, Dietrich Lehmann, Pedro Valdes Sosa, Elisabeth Braeker, Horst Kleinlogel, Robert Isenhart, and E Roy John (2002). “Millisecond by millisecond, year by year: normative EEG microstates and developmental stages”. In: Neuroimage 16.1, pp. 41–48.

Kullback, Solomon (1959). Information theory and statistics. Mineola, NY: Dover Publications, Inc.

Lehmann, D (1993). “Psychiatry and Microstates of the Brain’s Electric Field: Towards the “Atoms of Thought and Emotion”“. In: Imaging of the Brain in Psychiatry and Related Fields. Springer, p. 215– 222.

Lehmann, Dietrich, Pascal L Faber, Silvana Galderisi, Werner M Herrmann, Toshihiko Kinoshita, Martha Koukkou, Armida Mucci, Roberto D Pascual-Marqui, Naomi Saito, Jiri Wackermann, et al. (2005). “EEG microstate duration and syntax in acute, medication-naive, first-episode schizophrenia: a multi-center study”. In: Psychiatry Research: Neuroimaging 138.2, pp. 141–156.

Lehmann, Dietrich, Hisaki Ozaki, and Ivan Pál (1987). “EEG alpha map series: brain micro-states by space-oriented adaptive segmentation”. In: Electroencephalography and clinical neurophysiology 67.3, pp. 271–288.

Lehmann, Dietrich, Roberto D Pascual-Marqui, Werner K Strik, and Thomas Koenig (2010). “Core networks for visual-concrete and abstract thought content: a brain electric microstate analysis”. In: Neuroimage 49.1, pp. 1073–1079.

Lehmann, Dietrich and Wolfgang Skrandies (1980). “Reference-free identification of components of checkerboard-evoked multichannel potential fields”. In: Electroencephalography and clinical neurophysiology 48.6, pp. 609–621.

Lehmann, Dietrich, Werner Konrad Strik, Barbara Henggeler Thomas König, and Martha Koukkou (1998). “Brain electric microstates and momentary conscious mind states as building blocks of spontaneous thinking: I. Visual imagery and abstract thoughts”. In: International Journal of Psychophysiology 29.1, pp. 1–11.

Levin, David A and Yuval Peres (2006). Markov chains and mixing times. Vol. 107. Providence, RI: American Mathematical Soc.

Lizier, Joseph T, Mikhail Prokopenko, and Albert Y Zomaya (2012). “Local measures of information storage in complex distributed computation”. In: Information Sciences 208, pp. 39–54.

Lloyd, Stuart (1982). “Least squares quantization in PCM”. In: IEEE transactions on information theory 28.2, pp. 129–137.

Lütkepohl, Helmut (2005). New introduction to multiple time series analysis. Springer Science & Business Media.

Murray, Micah M, Denis Brunet, and Christoph M Michel (2008). “Topographic ERP analyses: a step-by-step tutorial review”. In: Brain topography 20.4, pp. 249–264.

Musso, Francesco, Jürgen Brinkmeyer, Arian Mobascher, Tracy Warbrick, and Georg Winterer (2010). “Spontaneous brain activity and EEG microstates. A novel EEG/fMRI analysis approach to explore resting-state networks”. In: Neuroimage 52.4, pp. 1149–1161.

Paluš, Milan (2007). “From nonlinearity to causality: statistical testing and inference of physical mechanisms underlying complex dynamics”. In: Contemporary physics 48.6, pp. 307–348.

Pascual-Marqui, Roberto D, Christoph M Michel, and Dietrich Lehmann (1995). “Segmentation of brain electrical activity into microstates: model estimation and validation”. In: IEEE Transactions on Biomedical Engineering 42.7, pp. 658–665.

Pedregosa, F., G. Varoquaux, A. Gramfort, V. Michel, B. Thirion, O. Grisel, M. Blondel, P. Prettenhofer, R. Weiss, V. Dubourg, J. Vanderplas, A. Passos, D. Cournapeau, M. Brucher, M. Perrot, and E. Duchesnay (2011). “Scikit-learn: Machine Learning in Python”. In: Journal of Machine Learning Research 12, pp. 2825–2830.

Ponce-Alvarez Adrián, Adrien Jouary, Martin Privat, Gustavo Deco, and Germán Sumbre (2018). “Whole-brain neuronal activity displays crackling noise dynamics”. In: Neuron 100.6, pp. 1446– 1459.

Rezek, Iead and Stephen J Roberts (2002). “Ensemble hidden markov models for biosignal analysis”. In: 2002 14th International Conference on Digital Signal Processing Proceedings. DSP 2002 (Cat. No. 02TH8628). Vol. 1. IEEE, pp. 387–391.

Rieger, Kathryn, Laura Diaz Hernandez, Anja Baenninger, and Thomas Koenig (2016). “15 years of microstate research in schizophrenia–where are we? A meta-analysis”. In: Frontiers in psychiatry 7, xp. 22.

Roweis, Sam T and Lawrence K Saul (2000). “Nonlinear dimensionality reduction by locally linear embedding”. In: science 290.5500, pp. 2323–2326.

Rué-Queralt, Joan, Angus Stevner, Enzo Tagliazucchi, Helmut Laufs, Morten L Kringelbach, Gustavo Deco, and Selen Atasoy (2021). “Decoding brain states on the intrinsic manifold of human brain dynamics across wakefulness and sleep”. In: Communications Biology 4.1, p. 854.

Rukat, Tammo, Adam Baker, Andrew Quinn, and Mark Woolrich (2016). “Resting state brain networks from EEG: hidden Markov states vs. classical microstates”. In: arXiv, p. 1606.02344.

Schlegel, Felix, Dietrich Lehmann, Pascal L Faber, Patricia Milz, and Lorena RR Gianotti (2012). “EEG microstates during resting represent personality differences”. In: Brain topography 25.1, pp. 20– 26.

Shine, James M, Michael Breakspear, Peter T Bell, Kaylena A Ehgoetz Martens, Richard Shine, Oluwasanmi Koyejo, Olaf Sporns, and Russell A Poldrack (2019). “Human cognition involves the dynamic inte-gration of neural activity and neuromodulatory systems”. In: Nature neuroscience 22.2, pp. 289– 296.

Skrandies, Wolfgang (1989). “Data reduction of multichannel fields: global field power and principal component analysis”. In: Brain topography 2.1, pp. 73–80.

Spencer, Kevin M, Joseph Dien, and Emanuel Donchin (1999). “A componential analysis of the ERP elicited by novel events using a dense electrode array”. In: Psychophysiology 36.3, pp. 409–414.

Spencer, Kevin M(2001). “Spatiotemporal analysis of the late ERP responses to deviant stimuli”. In: Psychophysiology 38.2, pp. 343–358.

Stevens, Andreas and Tilo Kircher (1998). “Cognitive decline unlike normal aging is associated with alterations of EEG temporo-spatial characteristics”. In: European archives of psychiatry and clinical neuroscience 248.5, pp. 259–266.

Theiler, James, Stephen Eubank, André Longtin, Bryan Galdrikian, and J Doyne Farmer (1992). “Testing for nonlinearity in time series: the method of surrogate data”. In: Physica D: Nonlinear Phenomena 58.1-4, pp. 77–94.

Tognoli, Emmanuelle and JA Scott Kelso (2014). “The metastable brain”. In: Neuron 81.1, pp. 35–48.

von Wegner, Frederic, Paul Knaut, and Helmut Laufs (2018). “EEG Microstate Sequences From Different Clustering Algorithms Are Information-Theoretically Invariant”. In: Frontiers in Computational Neuroscience 12, p. 70. ISSN: 1662-5188.

Wegner, Frederic von, Enzo Tagliazucchi, and Helmut Laufs (2017). “Information-theoretical analysis of resting state EEG microstate sequences-non-Markovianity, non-stationarity and periodicities”. In: Neuroimage 158, pp. 99–111.

Wilkinson, Mark D, Michel Dumontier, IJsbrand Jan Aalbersberg, Gabrielle Appleton, Myles Axton, Arie Baak, Niklas Blomberg, Jan-Willem Boiten, Luiz Bonino da Silva Santos, Philip E Bourne, et al. (2016). “The FAIR Guiding Principles for scientific data management and stewardship”. In: Scientific data 3.1, pp. 1–9.

Yuan, Han, Vadim Zotev, Raquel Phillips, Wayne C Drevets, and Jerzy Bodurka (2012). “Spatiotemporal dynamics of the brain at rest—exploring EEG microstates as electrophysiological signatures of BOLD resting state networks”. In: Neuroimage 60.4, pp. 2062–2072.

